# Corticosteroid resistance is predetermined by early immune response dynamics at acute Graft-versus-Host disease onset

**DOI:** 10.1101/2025.01.12.632608

**Authors:** Sophie Le Grand, Yannick Marie, Delphine Bouteiller, Émilie Robert, Régis Peffault de Latour, Gérard Socié, Nicolas Vallet, David Michonneau

## Abstract

Steroid-resistant acute graft versus host disease (SR-aGVHD) is the leading life-threatening complication following allogeneic hematopoietic stem cell transplantation. Novel therapeutics development is impeded by scares knowledge on biological pathways leading to steroid resistance at time of aGVHD diagnosis. To gain insight into our understanding on circulating immune cell subsets and functions at time of aGVHD, a single cell deep phenotyping and transcriptome analysis was performed on peripheral blood mononuclear cells from patients with aGVHD before steroid treatment or without aGVHD. We aimed at identifying biological patterns associated with steroid resistance at early onset of aGVHD. First, circulating immune cell subsets were associated with increased incidence of aGVHD, but not with steroid sensitivity. Then, pathway analysis and inferred ligand/receptor interactions revealed major functional divergences between steroid-sensitive (SS-) and SR-aGVHD, including enrichment of TNFα activation in SR-GVHD, as well as TNF/TNFR, CCL3, CCL4 and IL18 signaling, and decreased interferon α and γ signaling pathways, suggesting that steroid resistance in an intrinsic property of immune cells before any treatment. To go deeper into the understanding of mechanisms at play during SR-aGVHD, we modeled immune trajectories within CD8^+^ T cells and evidenced specific direct transition, from an early naive state to a highly activated one. By contrast, SS-aGVHD involved specific gene signatures across multiple intermediate differentiation stages during cell-to-cell transitions. These findings provide evidence that steroid resistance is driven by intrinsic mechanisms already present at the onset of alloimmune response, that may serve as potential new therapeutic targets.

## Introduction

Allogeneic hematopoietic stem cell transplantation (HSCT) is a major treatment for hematological malignancies. Its therapeutic effect is based on alloreactivity of donor T cells against tumor cells. However, donor T cells also react against healthy host cells, which may lead to graft-versus-host disease (GVHD). Acute GVHD (aGVHD) may involve skin, gut, or liver tissue. Its pathophysiology is initiated after tissue damage from high dose alkylating agent and radiation based-conditioning regimens leading to recognition of non-self-major and minor histocompatibility antigens (MHC and mHag(*1*)) and an alloreactive immune response, exacerbated by altered tissue repair mechanisms and microbiome dysbiosis(*2*). Despite first line treatment with prompt corticosteroid initiation(*3*, *4*), more than 40% of patients are steroid resistant (SR-aGVHD)(*5*, *6*). Steroid resistance is defined by a progressive disease after three days of standard dose of corticosteroids, an absence of clinical response after 7 days, an absence of complete remission after two to four weeks, or recurrence while steroids are stopped or tapered. Although ruxolitinib improves symptoms for 60% of SR-GVHD patients(*7*, *8*), their prognosis is very dismal(*3*, *7*, *9*).

While biomarkers were recently developed to improve aGVHD diagnosis and risk stratification(*10–14*), response to steroid is still unpredictable and biological mechanisms underlying steroid resistance remains only partly understood, even in the few experimental transplantation models of SR–GVHD described(*15–17*). In humans, specific whole gene expression analysis at diagnosis of lower intestinal acute SR-GVHD identified an increased repair-associated gene (amphiregulin and the aryl hydrocarbon receptor) and M2 macrophages pathways(*18*), suggesting that steroid resistance in not restricted to the T cells response as initially suspected. Indeed, neutrophils participate in tissue damage at GVHD onset and triggering, through reactive oxygen species production(*19*) and promote T cell expansion(*20*) and differentiation to Th17 cells(*21*, *22*). As corticosteroids increase neutrophils count and development, it has been hypothesized that corticosteroids may trigger a positive feedback loop between pathogenic T cells and neutrophils in SR-GVHD context(*23*).

Furthermore, neutrophil-predominant noncanonical inflammation pathways were recently identified in gut-SR-GVHD biopsies, notably ubiquitin specific peptidase 17–like family of genes(*24*). While gut microbiota’s role has been extensively studied in GVHD, it is not well understood in SR-GVHD. Butyrate, a short-chain-fatty-acid product from *Blautia* bacteria is associated with better long terms outcomes in aGVHD(*25–27*), but one study found a negative association between butyrate production and SR-GVHD, due to the ability of butyrate to impair human colonic stem cells from recovering epithelial monolayer(*28*). Other studies identified SR-GVHD biomarkers such as IL-22(*15*), but with controversial potential(*29*, *30*). Most of these studies rely on murine models(*15–17*, *23*, *31*), and there is currently no clear overview of immune subsets associated with steroid resistance in human GVHD.

We hypothesize that reshaping of immune system might be initiated early in alloimmune responses, therefore predisposing to corticosteroids resistance. To describe immune subsets and their respective function, cellular indexing of transcriptomes and epitopes sequencing (CITE-Seq) was performed on peripheral blood mononuclear cells (PBMC) samples collected at the onset of aGVHD before first line treatment, or in patient without GVHD at day 100 after HSCT. Whole landscape of circulating immune cells at early onset of aGVHD and its association with GVHD severity was uncovered. Deep analysis of transcriptomic profiles revealed that steroid resistance is an intrinsic property of the immune response, characterized by specific enriched biological pathways, specific cell-to-cell crosstalk through ligand-receptor interactions and a specific immune trajectory involving most immune subsets. Altogether, our results highlight that steroid-resistant GVHD is an early characteristic of allo-immune response before any immunosuppressive treatment and pave the way to more targeted first-line therapy to improve allogeneic HSCT outcomes.

## Results

### Cohort’s characteristics

Samples from 53 patients were retrieved from the prospective CRYOSTEM multicenter collection. PBMC were collected before corticosteroids initiation for patients diagnosed with aGVHD, and at day 100 post allo-HSCT for non-GVHD patients. Hematological malignancies, HSCT procedure, clinical and GVHD characteristics were balanced between groups, except for graft source, delay between sample and transplantation date (**Table S1)**. Among the 21 aGVHD patients, 16 (76%) were steroid sensitive (SS-GVHD) and 5 (24%) were steroid refractory (SR-GVHD), with a median delay of 24 days [13-96] and 17 days [11-70] between transplantation and sample, respectively (p=0.3) (**Figure 1A**). For validation and control purposes, 6 samples from healthy blood donor were used. After samples preparation for CITE-seq experiments, sequencing, quality control filtering and data processing acquisition, more than 300,000 single cells were available for immune profiling and transcriptome analysis (**Figure 1B**). Unsupervised clustering resulted in the identification of 40 immune cell subsets (**Figure 1C and S1**), characterized by their surface antigens expression (**Figure 1D**).

**Figure 1:**
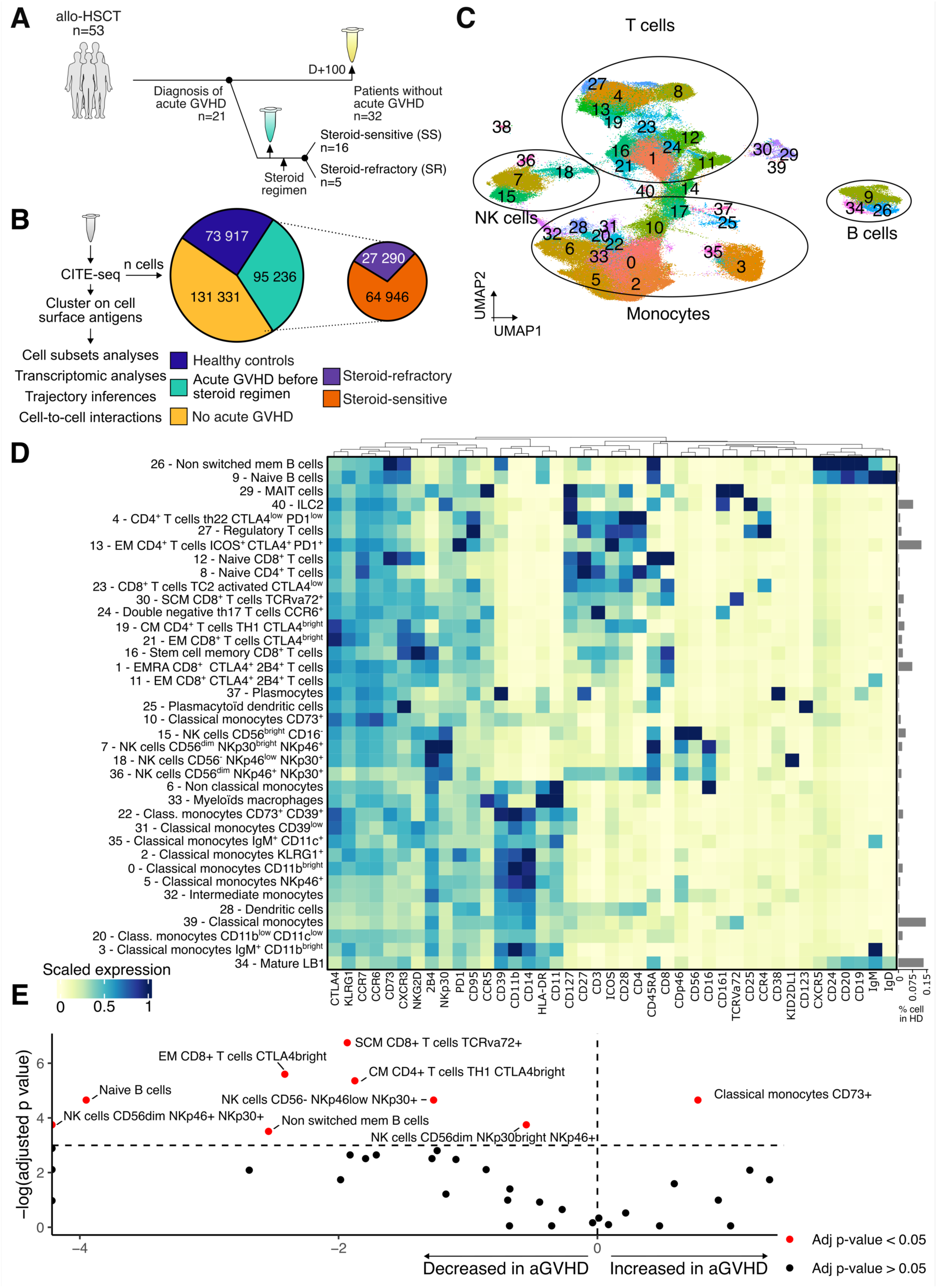
Overview of patient and cell distribution, cluster identification, and relative abundance of immune cell populations. **A:** Experimental design: HSCT patients (n=53) with (n=21) and without aGVHD (n=32) and healthy donor as control group (n=6) were analyzed. PBMC were collected at day 100 post HSCT for patients without aGVHD, and before steroid initiation for patients with aGVHD. **B:** Number of cells included in the analysis after CITE-Seq processing. **C:** Uniform Manifold Approximation and Projection (UMAP) after principal component analysis for immune cells clustering, according to surface markers. **D:** Heatmap depicting immune cluster’s identification through surface antigen expression. Three irrelevant populations were removed and are shown in **Figure S1**. **E**: Volcano plot exhibiting immune cells abundance in GVHD or non-GVHD condition. Y axis reflects que -log 10(adjusted p-value) from differential expression output, using non-parametric Mann-Whitney-Wilcoxon rank sum test. X axis reflects the log(ratio) of median abundance in aGVHD condition, over non-aGVHD condition. Abbreviation: CITE-Seq: Cellular indexing of transcriptomes and epitopes sequencing. aGVHD: acute graft versus host disease. HSCT: allogeneic hematopoietic stem cell transplantation. PBMC: Peripheral Blood Mononuclear Cells. SS-aGVHD: steroid sensitive acute graft versus host disease. SR-aGVHD: steroid resistant acute graft versus host disease.

### Immune signature and increased proliferation associate with aGVHD onset but not with response to steroids

Compared to non-aGVHD patients, classical monocytes CD73^+^ were significantly enriched in aGVHD patients, whereas NK and B cell subsets were significantly decreased (**Figure 1E**). To encompass the compositional structure of immune subsets frequencies, principal coordinates analysis (PCoA) was used. Four distinct immune clusters were identified, thereafter called immunotypes (**Figure 2A**). aGVHD patient’s distribution was significantly different among immunotypes (p=0.001, **Table S2**, **Figure 2B**), as well as the GVHD cumulative incidence (**Figure 2C**, Immunotype 4 GVHD incidence = 72% at day 100, *vs* immunotype 1 = 55%, *vs* immunotype 2 = 38%, *vs* immunotype 4 = 0%, global log-rank p=0.005). GVHD-free and relapse-free survival (GRFS) was significantly associated with immunotypes (p=0.0048, **Figure S2**), however there were no significant differences in non-relapse mortality, cumulative incidence of relapse or of chronic GVHD (**Figure S2**), and no association with aGVHD organ involvement (**Table S2**). Association with other clinical variables is depicted in **Figure S3**.

**Figure 2:**
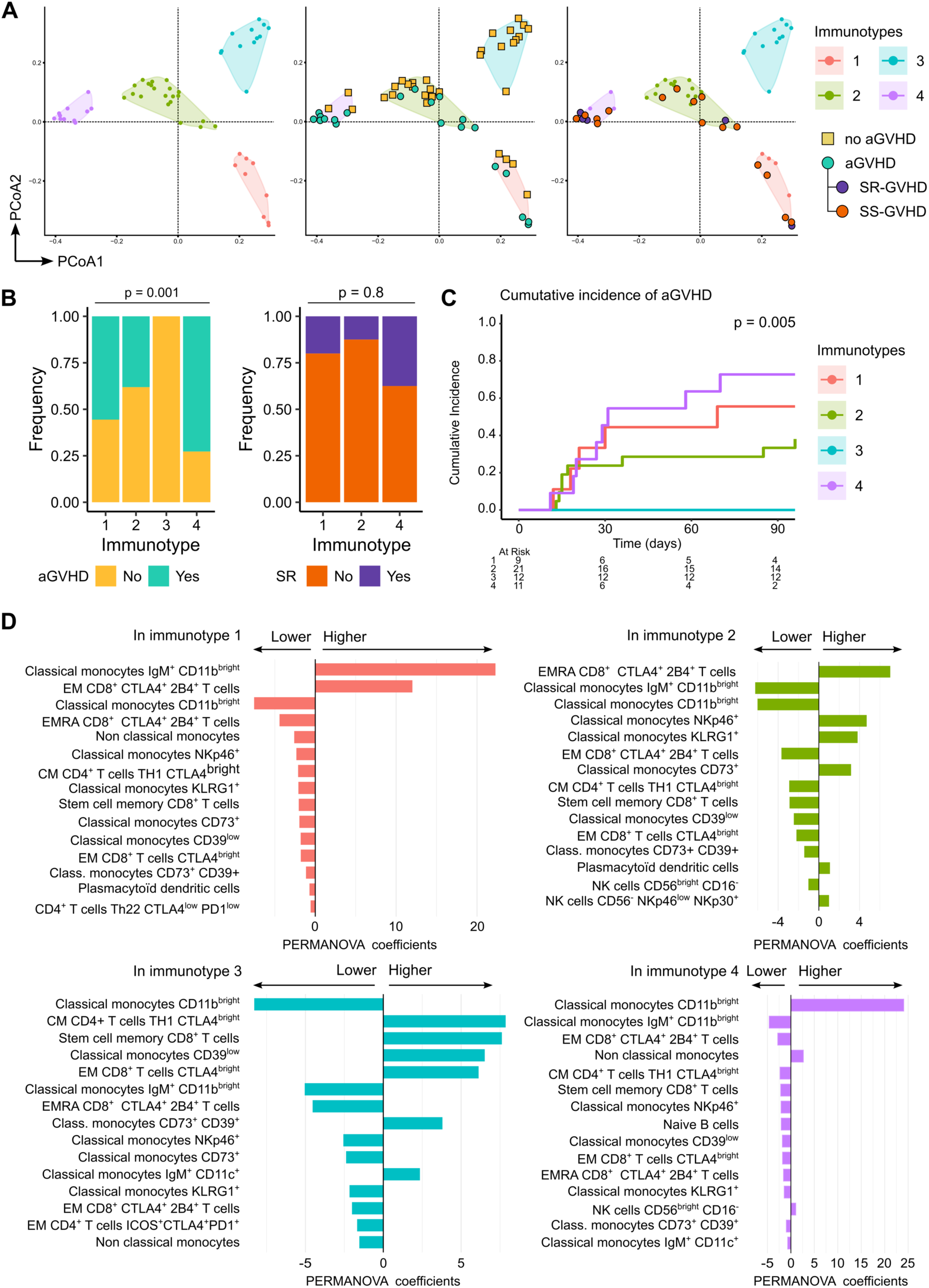
Immunotypes distinguished different immune patterns associated with aGVHD incidence. **A:** Principal coordinates analysis (PCoA) revealed 4 different immune clusters (immunotypes) according to cell population composition. **B:** Distribution of aGVHD patients among immunotypes (left panel) and corticosteroid sensitivity (right panel). Fisher test was used for statistical analysis. **C:** GVHD cumulative incidence with death as competing risk, estimated by Kaplan-Meir method, Log rank test was used for statistical analysis. Immunotype 4 cumulative incidence at day 100 = 72%, Immunotype 3 cumulative incidence at day 100 = 0%, Immunotype 2 cumulative incidence at day 100 = 38%, Immunotype 1 cumulative incidence at day 100 = 55%, log-rank p-value = 0.005. **D:** Bar plots showing top 15 contributing immune subsets to immunotypes. Contribution was determined with permutational multivariate analysis of variance (PERMANOVA).

Immunotype 1 and 4 included a large number of patients with aGVHD (**Figure 2B–C**) and were characterized by an overall reduction in immune cell heterogeneity, illustrated with an imbalanced immune subsets representation (**Figure 2D**). Especially, immunotypes 1 and 4 presented a high display of classical monocytes CD11b^bright^ (IgM^+^ or IgM^-^) compared to other immunotypes. Immunotype 2 and 3 were characterized by a balanced distribution across different immune cell populations, and a lower classical monocytes CD11b^bright^ cells representation. Remarkably, immunotype 3 was characterized by an absence of patients with aGVHD, a decrease in classical monocytes CD11b^bright^ and in other monocytes populations, whereas effector T CD4^+^ and CD8^+^ were predominantly increased.

To further investigate whether the over-representation may encompass a high cell division rate at time of aGVHD diagnosis, we determined cell cycle phase scoring within each immune cell population. Immune populations were predominantly in S phase (**Figure 3A**), reflecting increase cell proliferation at aGVHD onset.

**Figure 3:**
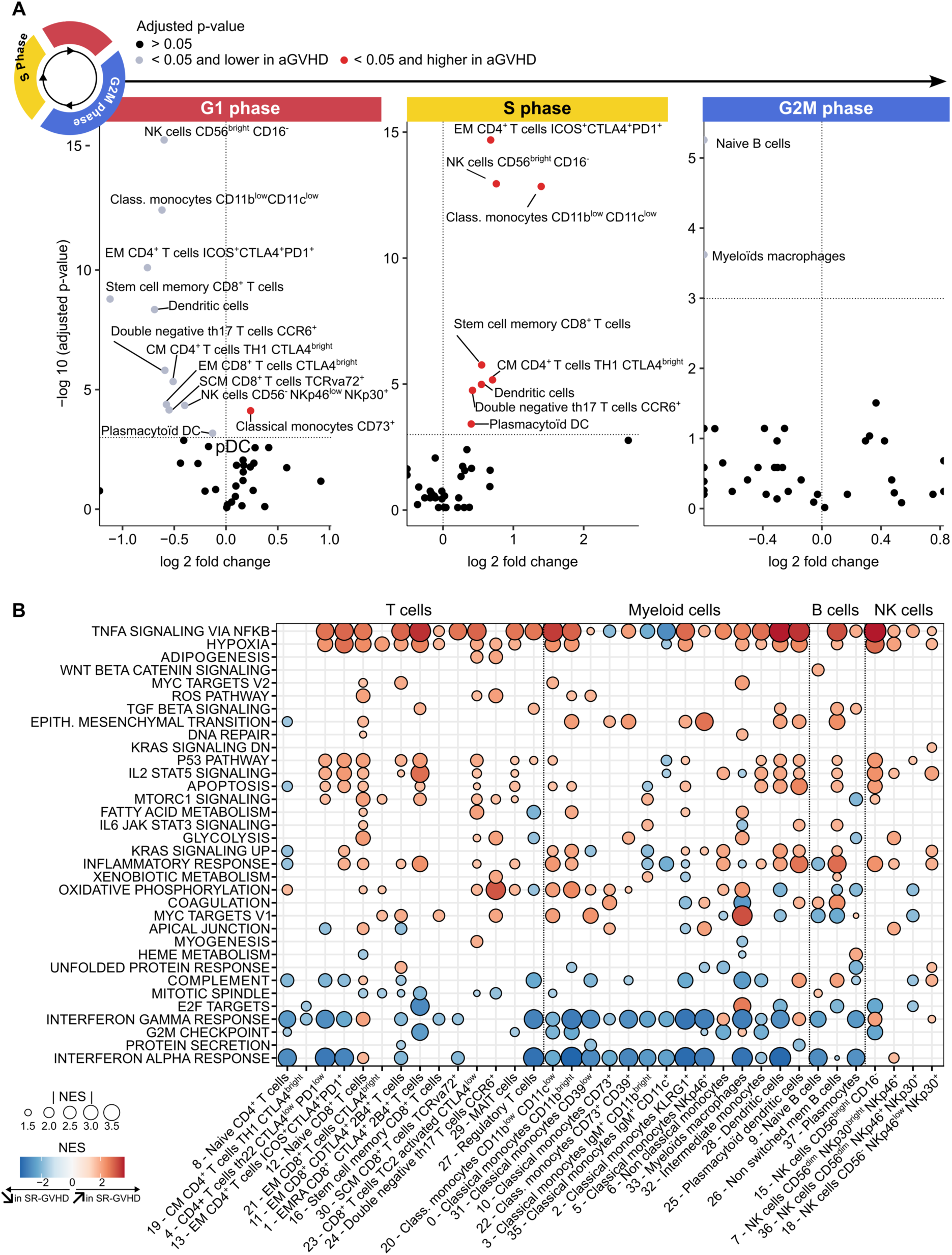
Cellular function analysis identified distinct features associated with SR-GVHD. **A:** Volcano plots exhibiting cell cycle analysis after cell cycle phases scoring. X axis represents log ratio of median number of cells in each phase in aGVHD patients over non GVHD patients. Y axis represents log adjusted p values. Immune population significantly increased or decreased are marked in red. Mann-Whitney-Wilcoxon tests were performed for statistical analysis, and p values were adjusted using Benjamini and Hochberg method. **B:** Pathways enrichment analysis comparing SR and SS-aGVHD, using gene set enrichment method and “hallmarks” gene reference data set. Only significant values with adjusted p-value < 0.05 are shown. NES: normalized enrichment score. SS-aGVHD: steroid sensitive acute graft versus host disease. SR-aGVHD: steroid resistant acute graft versus host disease.

No significant differences among immune cells abundances, immunotypes and cell-cycle scoring was observed with respect to corticosteroid resistance (**Figure 2B**, **Table S2, Figure S4 and S5**). In summary, immune cells subset clearly identified patients without GVHD but hardly discriminated SS and SR-GVHD. We thus hypothesized that cell intrinsic biological functions may predispose to SR-GVHD.

### Steroid resistance is associated with enrichment of TNFα and metabolic pathways at aGVHD onset

We first considered the biological pathways enriched at the onset of aGVHD by comparison with patients without GVHD (**Figure S6**). When considering aGVHD as a whole, we observe overactivation of IFNα and γ pathways in all immune cells, and a decrease in TNFα signaling, especially in CD8^+^ T cells and myeloid cells.

To study which biological process may correlate with subsequent SS- or SR-aGVHD, we then performed a second enrichment analysis in patients with SS- or SR-aGVHD. Compared to SS-aGVHD, inflammation pathways were enriched in SR-aGVHD patients within all immune cells, particularly TNFα and hypoxia pathways (**Figure 3B**). STAT5 and MTORC pathways were particularly upregulated in exhausted effector memory CD4^+^ and CD8^+^ T cells, in naïve CD8^+^ T cells and in dendritic immune cells. The IL6/JAK STAT pathway was also highly enriched in naive CD8^+^ T cells and classical CD11b^bright^ monocytes. Genes associated with metabolism pathways were overexpressed, especially oxidative phosphorylation in naïve CD4^+^ T cells, double negative T cells and myeloid cells. Conversely, there was a decreased activation of the IFNα and γ responses within all immune subsets, except NK cells.

### Cell to cell crosstalk during SR-GVHD specifically involves IL1, CCL3, CCL4, TNFα and IL18 signaling

To address whether cell biological processes involved in SR-aGVHD may be involved in cell-to-cell interactions leading to early steroid-resistance, we examined intercellular communication using a computational inferring method(*32*). When focusing on CD8^+^ T cells as receiver, 20 ligands were identified as prioritized ligands, and their expression on sender cells is shown in **Figure 4A**. IL1B, CCL3 and CCL4 ligands were particularly overexpressed in SR-aGVHD condition (**Figure 4B**). Regulatory potential of each ligand with predicted targets genes in shown in **Figure 4C**.

**Figure 4:**
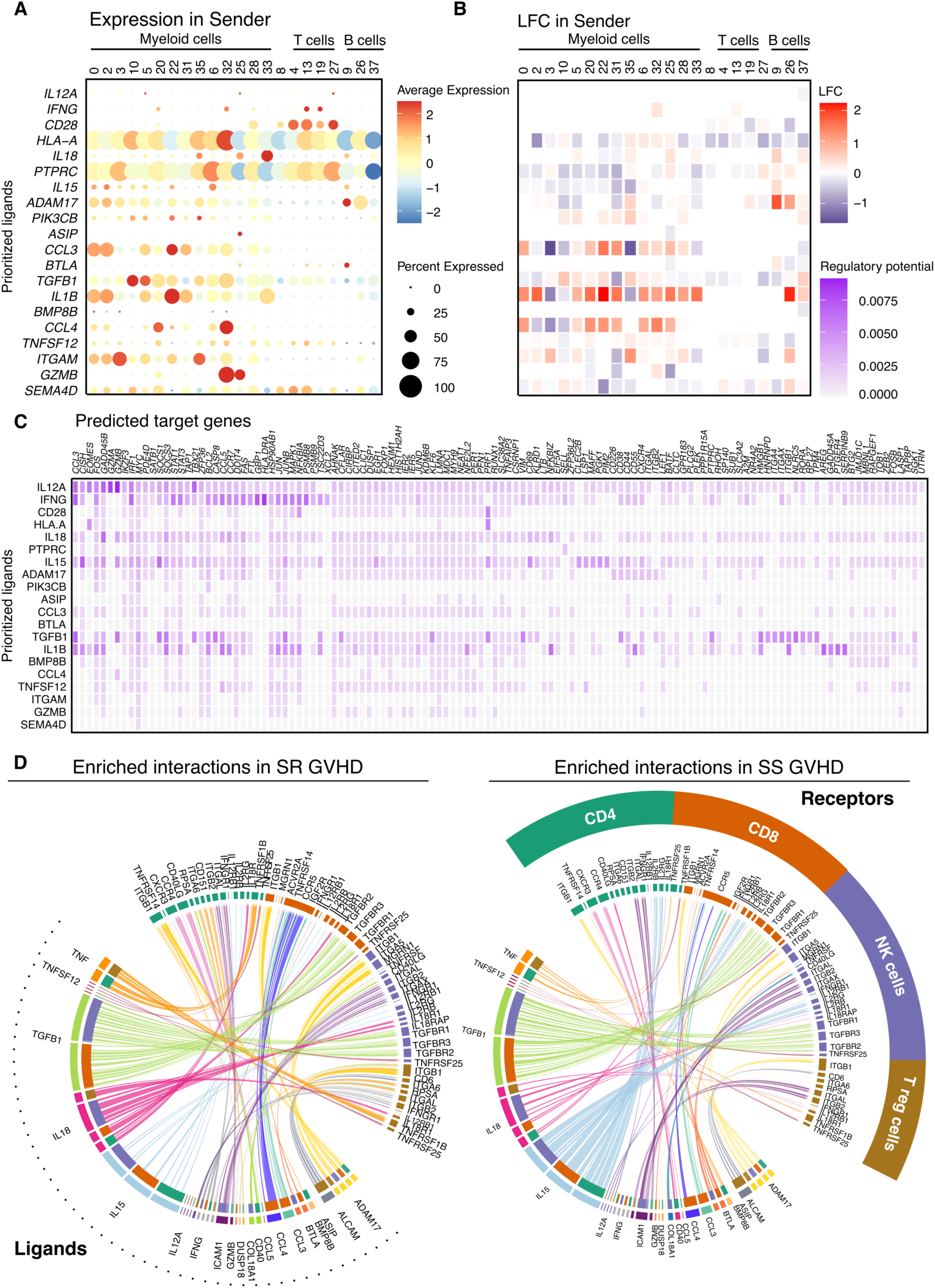
Cell-cell interactions with CD8+ T cells as receiver revealed specific patterns associated with SR-aGVHD. NicheNet Output when focusing on CD8+ T cells as receiver is depicted on panel A, B and C. **A:** Expression of top-ranked ligands on sender cells. Prioritized ligands identified are exhibited on Y axis, and sender cells on X axis. The color determines the average expression and the point size the percent of senders expressing the ligand. **B:** Relative expression of ligands on sender cells. Expression is exhibited in log fold change (red: increase expression in SR-aGVHD patients, Blue: decreased expression in SR-aGVHD patients). **C:** Ligand–target matrix exhibiting the regulatory potential of interactions. On the X axis is shown the active targets genes of the top-ranked ligands. Heatmap visualization showing which top-ranked ligands are predicted to have regulated the expression of which differentially expressed genes. **D:** Circos plots depicting interactions between senders and all receiver cells. Ligands are shown on the left side of the circos plot, and targets on the right side. Only *bona fide* interactions were selected (**Figure S9A**). Links exhibits positive log fold change (left panel, increased in SR-aGVHD patients) or negative log fold change (right panel, increased in SS-aGVHD patients) from “ligand differential expression heatmap” NicheNet’s output. The thickness of the link represents the numbers of sender populations involved in the interaction. SR-aGVHD: Steroid resistant acute graft versus host disease. SS-aGVHD: Steroid sensitive acute graft versus host disease.

Predicted ligands and targets interaction were then analyzed for CD4 regulatory T cells (**Figure S7A**), NK cells (**Figure S7B**), CD4^+^T cells (**Figure S8A**), and double negative T cells (**Figure S8B**). An overview of all these interactions is depicted as a circus plot (**Figure 4D**) based on biologically confirmed interactions previously designated as *bona fide* interactions (**Figure S9A**). TNF ligands-to-cell signaling appeared to be overexpressed in most sender cells in SR-aGVHD patients, as well as CCL3, CCL4 and IL18 signaling. ADAM17, which is involved in TNF activation(*33*), was overexpressed in all senders and interacted with TNFR, ITGAL and ITGB2 receptors. Interestingly, IFN/IFNR interactions was enhanced across all receivers in SR-aGVHD but absent in SS-aGVHD.

Conversely, when comparing patients with aGVHD to those without aGVHD, immune cells crosstalk appeared different to that disclosed in the SS- and SR–aGVHD setting (**Figure S9B**). Consistent with enrichment analysis, TNF/TNF-R interaction was decreased among DN T cells, NK cells and T regulatory lymphocytes. TGFB1/TGFBR interactions, which are considered as being associated with an immunoregulatory role, were significantly increased in patient with no aGVHD compared to aGVHD patients, especially within CD8^+^ T cells. BTLA/TNFRSF14 interaction is another pathway previously identified in animal models as a regulator of alloimmune response(*34*, *35*), and was overexpressed in patients without GVHD.

### Specific trajectories were associated with SR-GVHD

As biological processes and cell-to-cell interactions targeting CD8^+^ T cells were associated with SR-aGVHD, we hypothesized that immune cells subsets from SR-aGVHD and SS-aGVHD patients may exert specific CD8^+^ T cell trajectories. To address this hypothesis, trajectories within CD8^+^ T cell were defined on a new cell clustering based on mRNA expression (**Figure S10**). Two trajectories were identified for CD8^+^ T cell: (i) lineage 1 described in **Figure 5A-B** and (ii) lineage 2 described in **Figure S11**.

**Figure 5:**
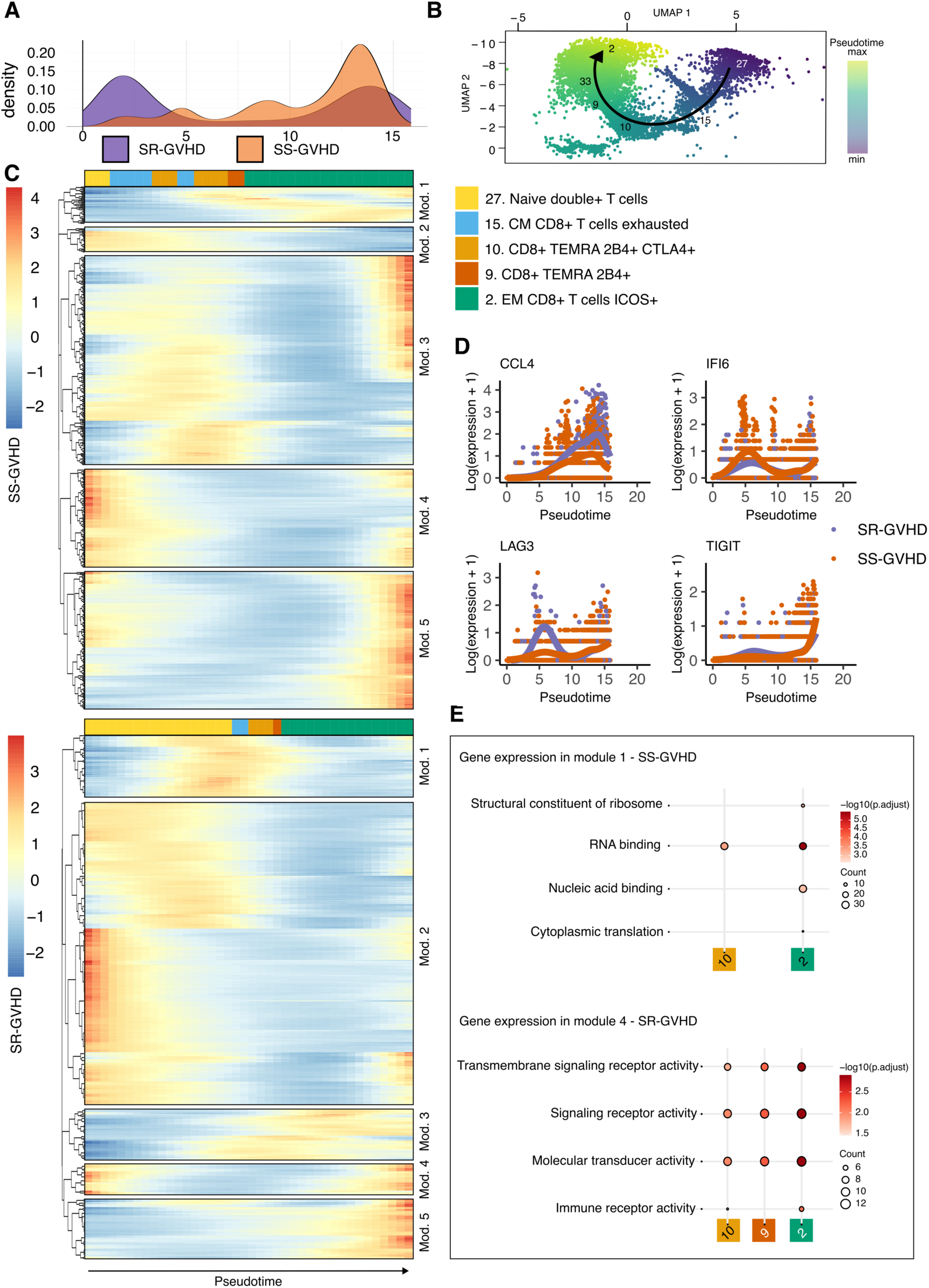
SR-aGVHD is associated with specific trajectory in CD8^+^ T cells. **A:** Density plot for number of cells according to lineage 1 pseudotime visualization. SR-aGVHD cell abundance was marked in low and high pseudotimes, while SS-aGVHD was associated with a continuous increase in cell abundance along differentiation. **B:** Lineage 1 trajectory on RNA clustering. **C:** Heatmap depicting gene expression among the top 500 most expressed genes in SS-aGVHD patient (top panel) and in SR-aGVHD patients (low panel), along pseudotime. The color at the top of each heatmap represents the immune population in which the gene is expressed. **D:** Expression along pseudotime of four of the top 100 genes, whose expression varied the most between the start and end points of a lineage. **E:** Over-representation analysis to identify pathways associated with genes expression from each module of the heatmaps. Only pathways associated with adjusted pval < 0.05 from “molecular functions” and “biological process” of the Gene Ontology knowledgbase(*63*, *64*) were selected for dot plots visualization. SR-aGVHD: Steroid resistant acute graft versus host disease. SS-aGVHD: Steroid sensitive acute graft versus host disease.

Frequency of cells in lineage 1 slightly increased over pseudotime in SS-aGVHD patients through multiple transition states starting from naïve double positive T cell (cluster 27) to effector memory CD8^+^ T cell expressing ICOS (cluster 2) **(Figure 5A-B).** By contrast, SR-aGVHD patients were characterized by a direct transition with very few cells in the transition subsets (cluster 15, 9 and 10), most immune subset being grouped in cluster 27 at early pseudotime or in cluster 2 in late pseudotime. The same pattern was observed in the second CD8^+^ T cell trajectory, with a transition from naïve double positive T cell (cluster 27) to CD8^+^ TEMRA cells expressing ICOS and PD1 (cluster 25) (**Figure S11A-B**).

To highlight different gene expression pattern in SS-aGVHD and SR-aGVHD we first studied individual genes whose expression varied significantly along pseudotime and across transition state (**Figure 5C, S11C, Table S3-S10**). Among the most significant genes involved in transition states, we identified CCL4, CCL5, IFI6, LAG3, and TIGIT (**Table S11 and S12**). SR-aGVHD patients exhibited earlier and higher CCL4 and LAG3 expression in pseudotime (**Figure 5D and S11D**). Consistently with pathway enrichment analysis, IFI6 expression, indicative of IFNα activation, was higher in SS-aGVHD patients, particularly in early-differentiated CD8^+^ T cells.

To study overall biological processes, we used hierarchical clustering to identify modules of correlated gene among the top 500 expressed gene along pseudotime lineages. This approach identified 5 modules of genes involved in cell-to-cell transition (**Figure 5C**). Over-representation analysis (ORA) from Gene Ontology knowledgebase within each cell subset highlighted the main biological pathways involved at each transition state (**Figure 5E**). SR-aGVHD was notably associated with an early enrichment of pathways associated with cell activation and high transcriptional and translational activity in naïve double positive T cell (**Figure S11E**). A detailed analysis of pathways enriched in each module revealed striking differences in the dynamics of T cell activation during SR-GVHD (trajectory 1 in **Figure S12 and S13,** and trajectory 2 in **Figure S14 and S15**).

Overall, these results reveal that even before starting corticosteroids, aGVHD is already characterized by different trajectories and pattern of gene activation between transition states, that can define a specific dynamic of alloimmune response during steroid resistance.

## Discussion

Despite recent advances in the management of aGVHD, steroid-resistant treatment remains a challenge. This study is the first to describe immune subsets and transcriptional profiles in human at time of aGVHD. Early identification of patients at risk of developing steroid resistance is crucial for rapid and tailored therapeutic intervention. Up to now, a comprehensive understanding of the circulating immune cells mechanisms associated with steroid resistance has not been addressed. Our results highlight that steroid resistance is characterized at the onset of aGVHD, by a specific transcriptomic profile that reveal biological pathways, cell-to-cell interactions and trajectories that differ from steroid sensitive patients.

Immunotypes and cell cycle scoring enabled the identification of specific patterns associated with aGVHD compared to non-aGVHD patients. Immunotype 4, characterized by poor immune cell diversity and by classical CD11^+^ monocytes abundance was associated with worsened aGVHD free survival. This observation is consistent with previous studies identifying monocytes and macrophages as fully parts of the immune response at aGVHD onset (*18*, *36–39*).

However, cell subsets distribution did not allow to distinguish SS- and SR-aGVHD. When focusing on immune cells functions rather than distinct phenotypes, we found specific patterns associated with SR-GVHD, especially higher TNFα pathway activation. This was strengthened with inferred interaction analyses which revealed an increased TNF/TNFR signaling, notably in CD8^+^ T cells. Although this *in silico* prediction would need to be confirmed *in vivo*, these results might suggest that early anti-TNF therapies could be beneficial in patients with the corticosteroid resistance profile we described herein. This is supported by a previous clinical phase 2 trial testing a combined therapy of anti-TNF and corticosteroids, with increased response rate (69% vs 33%; P < 0.001) without increased infections risk(*40*).

Our results suggest that the alloimmune response responsible for GVHD is already different at its initiation in patients who will be steroid-resistant. Overall, the integration of our data shows that steroid resistance could result from the cytokine production profile by myeloid populations, especially monocytes producing CCL3, CCL4, IL1, and IL18, which stimulate the activation of naive CD8^+^ T cells and induce direct differentiation into effector T cells (**Figure 6**). These results are consistent with published data on the role of myeloid populations(*19*, *20*) and inflammasome activation(*41*) in the initiation of acute GVHD in mice, or of M2 macrophage in the gut of patients with SR-GVHD(*18*). Additionally, previous data had also suggested that the effect of IL-18 differs depending on whether GVHD is mainly mediated by CD8^+^ or CD4^+^ T cells in murine models(*42*). Our data suggest that, in humans, this effector phase depends on the activation profile of CD8^+^ T cells, and that IL-18 may worsen the severity of CD8-mediated GVHD by increasing steroid resistance. IL18 is a proinflammatory cytokine which can promote interferon γ production(*43*), and its serum level in post-HSCT patients was associated with aGVHD severity (r = 0.861, P < 0.001)(*44*). Furthermore, our results highlight the key role of TNFα production by myeloid cells in SR-GVHD on CD4+ T activation that could amplify the alloimmune response. These results confirm data obtained in animal models and show that myeloid subsets may not only participate in the initiation of GVHD(*19*) but that their response profile could already initiate the first mechanisms of steroid resistance.

**Figure 6:**
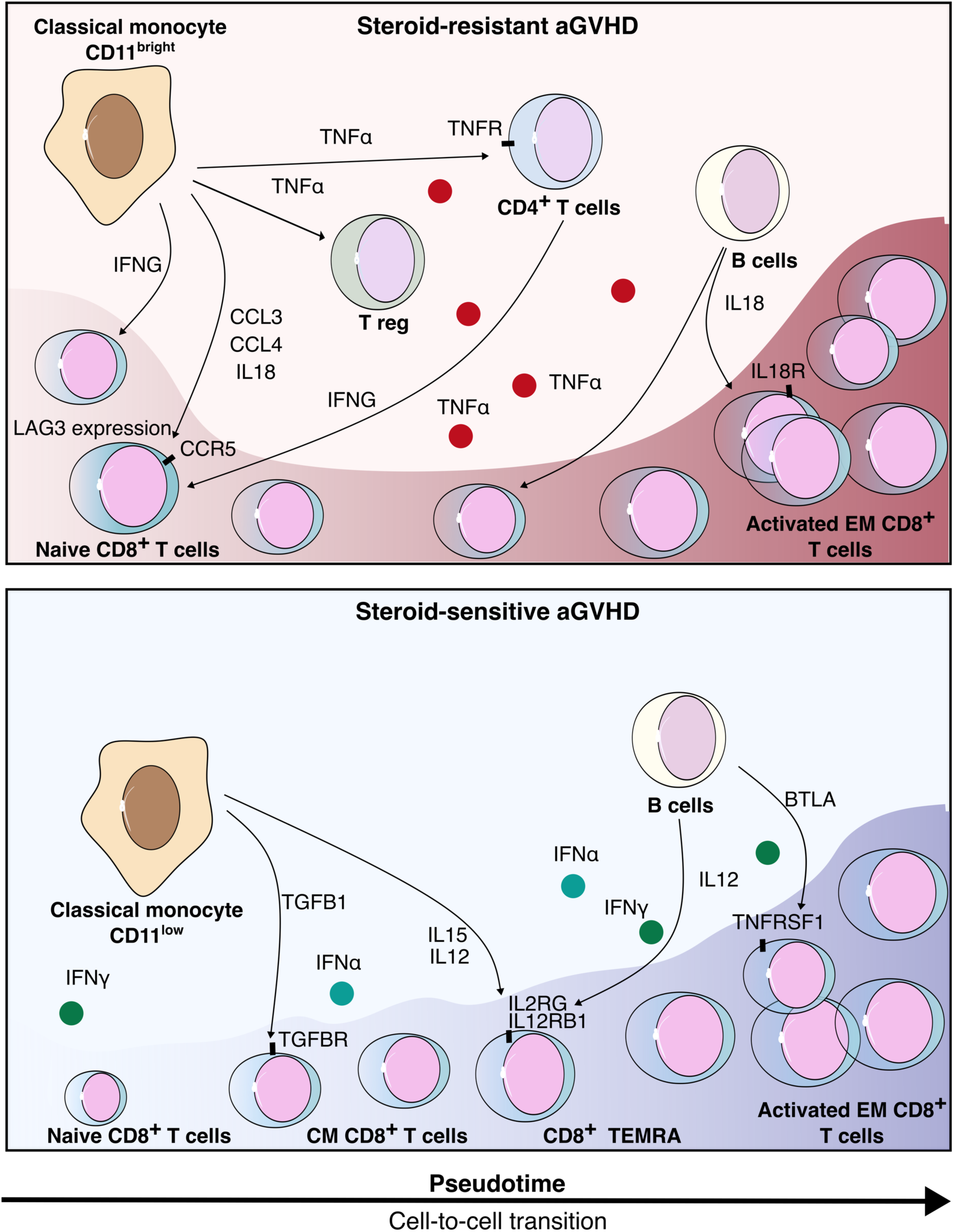
Steroid resistance is driven by biological pathways and specific networks of interaction. Steroid-resistant aGVHD results of cytokine production by myeloid populations, especially classical monocytes producing CCL3, CCL4, and IL18, which stimulate the activation of naive CD8^+^ T cells and induce direct differentiation into effector T cells. By contrast, steroid-sensitive aGVHD is mainly driven by IL15 and IL12 production from classical monocytes and B cells, and multiple transition states, from naïve CD8^+^ T cells to activated EM CD8^+^ T cells. aGVHD: acute graft versus host disease, CM CD8^+^ T cells: Central Memory CD8^+^ T cells, EM CD8^+^ T cells: Effector Memory CD8^+^ T cells, TEMRA CD8^+^ T cells: Terminally differenciated Effector Memory CD8^+^ T cells

Interferon have been described for their crucial role in GVHD development(*45–47*). We found that IFNα and γ pathways were enriched at time of aGVHD and before corticosteroid treatment. However, SR-aGVHD was associated with lower activation of IFNα and γ pathways in most immune subsets, except for naïve CD8 T cells and non-classical monocytes. This could explain why in SR-aGVHD, INFG/IFNGR interactions increased cell-to-cell crosstalk between CD8+ T cells, monocytes and CD4+ T cells. In addition, it has been suggested previously that IFNγ production could contribute to increase the severity of digestive GVHD(*48*, *49*) and that IFNλ blockade could improve gut GVHD severity(*50*).

Finally, trajectory inferences highlighted specific layouts associated with SR-aGVHD. Cells were particularly abundant in low and high pseudotime in SR-GVHD patients, while there was a progressive increase along pseudotime in SS-GVHD patients **(Figure 6)**. It suggests that cells in SR-GVHD patients quickly differentiate to terminally activated states. This was consistent with, higher and earlier gene expression of LAG3 and TIGIT in SR-GVHD patients. Consistent with results from cell-cell interactions analysis, CCL4 and CCL5 were expressed at higher levels and earlier in SR-GVHD patients, as also described in animal model(*51*).

Because samples were collected before cortico-steroid initiation, our work sheds light on early signature pathways leading to steroid resistance. These pathways appeared distinct from those involved in the general inflammatory response of acute GVHD and could be targeted as early as steroid initiation. Transcriptomic analysis offered a powerful approach to unravel immune cells functionality, combined with precise phenotypic population identification, allowing for the identification of cellular heterogeneity in SR-GVHD immune response. The complexity of the immune response in the context of GVHD was already investigated at the single cell level, through T cell subset analysis following TCR clonotype mapping(*52*). This elegant approach, revealed that each patient possessed a highly intricate repertoire, with different tissues exhibiting distinct clonal populations, varying degrees of repertoire overlap between sites, and diverse numbers of dominant clones shared across multiple tissues (*52*). More recently, a similar approach highlighted the impact of the tissue microenvironment on maintaining T-cell activation and driving divergent immune responses in peripheral organs (*53*). Our results rely on PBMCs samples and hence, provide a complementary perspective on acute GVHD, that uncover biological pathways associated with steroid resistance, irrespectively of targeted organs. Studies based directly on blood samples are essential in clinical practice, as they provide an easy access to potential predictors that could guide therapeutic management.

Our study offers new insights into the immune complexity, presenting a broader perspective of the immune response. Since our samples were all collected at onset of GVHD, the sampling time varied between patients, which could impact the immune response, particularly cytokine secretion. Therefore, while our study underscores the need for validation in a larger cohort with careful matching of sampling times, it could also serve as a robust basis for future studies involving a broader patient population. Furthermore, the use of models would require additional validation *in vivo*. Finally, additional studies focusing on immune infiltrates in target organs of GVHD would certainly help to complete our understanding of acute GVHD pathophysiology, and especially of mechanisms associated with corticosteroid resistance.

Overall, these findings provide evidence that corticosteroid resistance is an intrinsic mechanism already present at the onset of alloimmune response, highlighting specific patterns that may serve as potential new therapeutic targets.

## Materials and Methods

### Cohort

This study was conducted with annotated samples from patients and healthy donors provided by the CRYOSTEM Consortium and the SFGM-TC (Société Francophone de Greffe de Moelle et de Thérapie Cellulaire). Patients were included in the CRYOSTEM cohort after validation by the scientific committee (study number CS 18-01). CRYOSTEM cohort and collection were approved by IRB Sud-Méditerranée 1 (reference number DC-2014-2312 and DC-2019-3425) and the Commission Nationale de l’Informatique et des Libertés for data protection (reference number nz70243374i n°912120). All patients gave their written consent for clinical research and sample collection. This non-interventional research study with no additional clinical procedure was carried out in accordance with the Declaration of Helsinki. Data analyses were carried out using an anonymized database. Healthy donor’s PBMC were isolated from residual blood after apheresis provided by Etablissement Français du Sang (18/EFS/032).

### CITE Seq experiment

After thawing cells in a 50% fetal calf serum (FCS)/RPMI medium, 10x Genomics Feature Barcoding recommendations were followed. Cells were washed twice using a 10 mL solution of the same media. 500 000 cells were transferred in a 0.04% PBS/bovine serum albumin (BSA) medium and washed. Cells were then suspended in a 1% PBS/BSA solution before incubation with oligonucleotide-coupled antibodies TotalSeq-B (panel described in **Table S13**) for 60 min on ice. Cells were washed 4 times (4°C, RCF 300, 5 minutes) and suspended to obtain a concentration of 1 000 cells/µL. 10 000 cells were subjected in Gel Bead Emulsion using Chromium 10x Genomics controller, according to manufacturer guidelines. Chromium™ Next GEM Single Cell 3’ Kit v3.1 (10X Genomics, cat. 1000268) and 3’ Feature Barcode Kit (10X Genomics, cat. 1000262) were used to prepare reagents.

To perform single cell RNA sequencing after cDNA amplification, the concentration of each sample was measured using Tapestation 2200 (Agilent). To prepare the cDNA libraries for 10x Genomics Chromium controller, we used the single-cell 3′ v3.1 kit (Dual Index) with Feature Barcode technology for Cell Surface Protein, following manufacturer guidelines. QC libraries were performed using Tapestation 2200 (Agilent). Libraries were equimolar pooled to obtain at least 20 000 read pairs per cell and 5 000 read pairs for feature barcode part after sequencing on Illumina Novaseq 6000 (100 cycles cartridge). The input number of cells was estimated at 10 000 cells/samples. FASTQ files were obtained with bcl2fastq (Illumina).

### Data quality control

Alignment, filtering, barcode counting, UMI counting were performed with count function from 10x Genomics Cell Ranger (version 6.0) using GRCh38 genome assembly as reference data. Then aggr function was used to aggregate results without normalization. A R environment (version 4.3.1) was subsequently used for the following analyses. Briefly, using Seurat R package(*54*), dead cells were excluded if mitochondrial RNA was above 18%; singlet were identified as droplets with RNA feature between 200 and 4 700; RNA and cell surface antigen counts were normalized and scaled using Seurat R package normalization and scaling function for linear transformation. 2 000 highly variable RNA features were identified.

### Immune cell clustering

Using a principal component analysis method on the scaled data, cells were clustered according to cell surface antigen expression. We used UMAP for cluster visualization. Seurat R package was then used to calculate cell cycle phase based on canonical transcripts, and regressing these out of the data.

### Enrichment pathways analysis

Fast pre-ranked gene set enrichment analysis was performed using FGSEA package(*55*) with “hallmarks” gene reference data set from Molecular signature database(*56*). Pathways were compared between GVHD and non-GVHD patients, and between SR-GVHD and SS-GVHD patients. Gene expression comparison between GVHD and non GVHD patients, and between SS- and SR-GVHD patients was provided by differential expression analysis. Enrichment scores associated with adjusted p-value < 0.05 were selected for dot plot visualization.

### NicheNet Analysis

Ligand-receptors interaction was computed using NicheNet package(*32*). Sender cells were identified as myeloid cells, B cells, CD4^+^ conventional T cells and regulatory T cells. Receiver cells were identified as CD4^+^ and CD8^+^ T lymphocytes, NK cells and regulatory T cells. Three pooled human gene expression data of interacting cells(*57–59*) were used as input and combined with a prior model that integrates existing knowledge on ligand-to-target signaling paths to perform ligand activity analyses. Circos plot visualization integrated *bona fide* interactions of each receiver, and increased or decreased interactions were integrated from “ligand differential expression heatmap” NicheNet output.

### Trajectory inferences

Trajectory were based on RNA expression clustering (**Figure S10**). SingleCellExperiment object was subset by immune populations (CD8+ T cells, CD4+ T cells, NK cells, B cells and myeloid cells) before pseudotime calculation using slingshot package(*60*). TradSeq package was then use for downstream analysis(*61*). Briefly, associationTest R function evidenced genes whose expression is associated with pseudotime. Since pseudotime is a continuous value, it was divided into 50 categories for graphical representation with heatmap. Only the 500 most expressed gene in SS- and SR-GVHD were selected for heatmap visualization (**Table S3-S6**). Each point on X axis corresponded to a cell cluster with similar pseudotimes. The immune populations represented on top of the heatmap corresponded to the immune cell population most abundantly represented within the cell cluster (**Table S7-S10**). Gene markers specific to low pseudotime or high pseudotime were found thanks to starVsEndTest command. Expression along pseudotime in each condition was plotted for four genes from the top 100 starVsEndTest output **(Figure 5, Tables S11 and S12)**. To explore the pathways associated with gene expression of the top 500 most expressed genes along pseudotime from each of the 5 modules of the heatmap, we used over-representation analysis using enrichGO command from clusterProfiler R package(*62*). Only pathways associated with adjusted pval < 0.05 from “molecular functions” and “biological process” of the Gene Ontology knowledgbase(*63*, *64*) were selected for dot plots visualization.

## Acknowledgments

The authors thank the patients who agreed to participate in this study by providing their blood samples. We also thank the iGenSeq platform team at ICM for sequencing the PBMC samples. We are grateful to the CRYOSTEM Consortium and the SFGM-TC for collecting patient samples and clinical data.

## Author contributions

Conceptualization: SLG, DM; Methodology: SLG, NV, DM; Samples processing: SLG, DB, YM, EB; Investigation: SLG, NV, DM; Visualization: SLG, GS, RPD, NV, DM; Funding acquisition: DM; Project administration: EB, DM, GS, RPD; Supervision: NV, DM; Writing – original draft: SLG, NV, DM; Draft revision: SLG, YM, DB, EB, RPF, GS, NV, DM.

## Data availability

Raw data are available on GEO public repository: GSE229733. Source code to reproduce the analyses, figures, and tables described in this manuscript are provided in a Git repository: https://gitlab.com/SophieLG/gvh_transcriptomic and long term preserved in Software Heritage: https://archive.softwareheritage.org/browse/origin/directory/?origin_url=https://gitlab.com/SophieLG/gvh_transcriptomic&timestamp=2024-12-09T14:03:56.753000%2B00:00

