## Supplementary material for "Corticosteroid resistance is predetermined by early immune response dynamics at acute Graft-versus-Host disease onset"

#### Supplementary Materials

### Supplementary figures

#### Figure S1

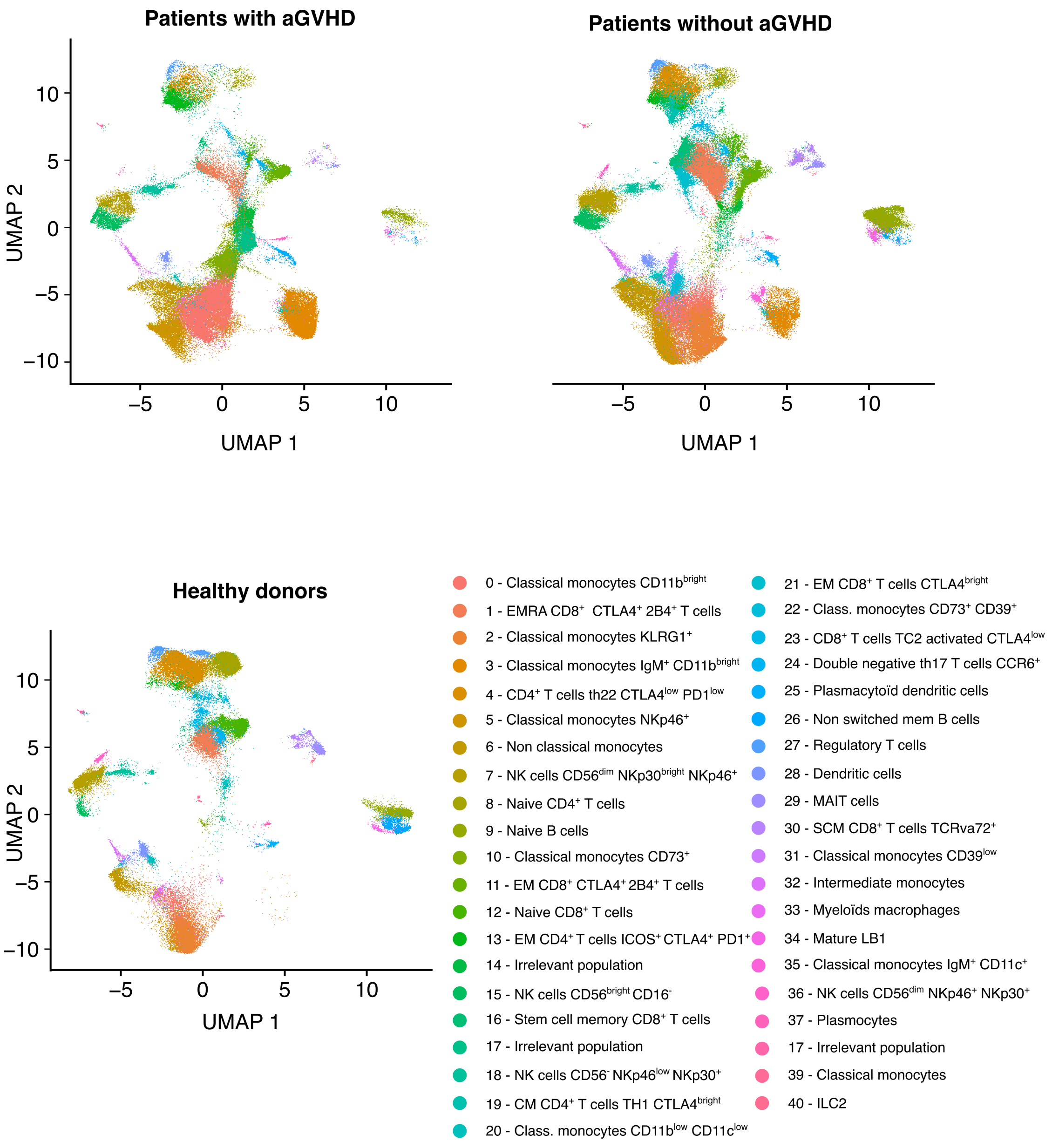


**Fig. S1: Immune populations in each group of patients:**

HSCT: hematopoietic Stem Cell Transplantation. GVHD: Graft Versus Host Disease.

**Figure S2**

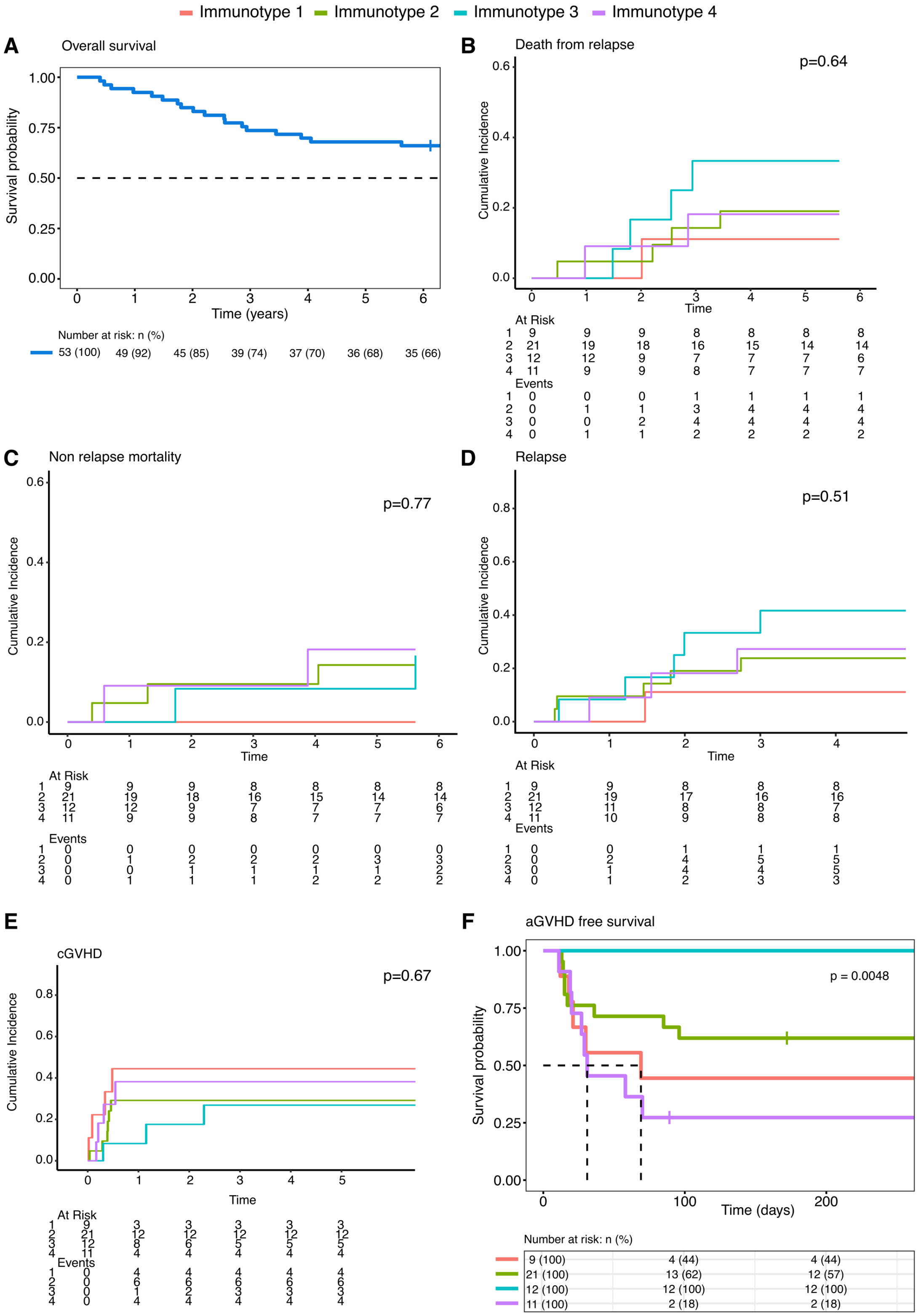


**Fig. S2: Immunotype survival analysis:**

Survival analyses according to immunotypes

**Figure S3**

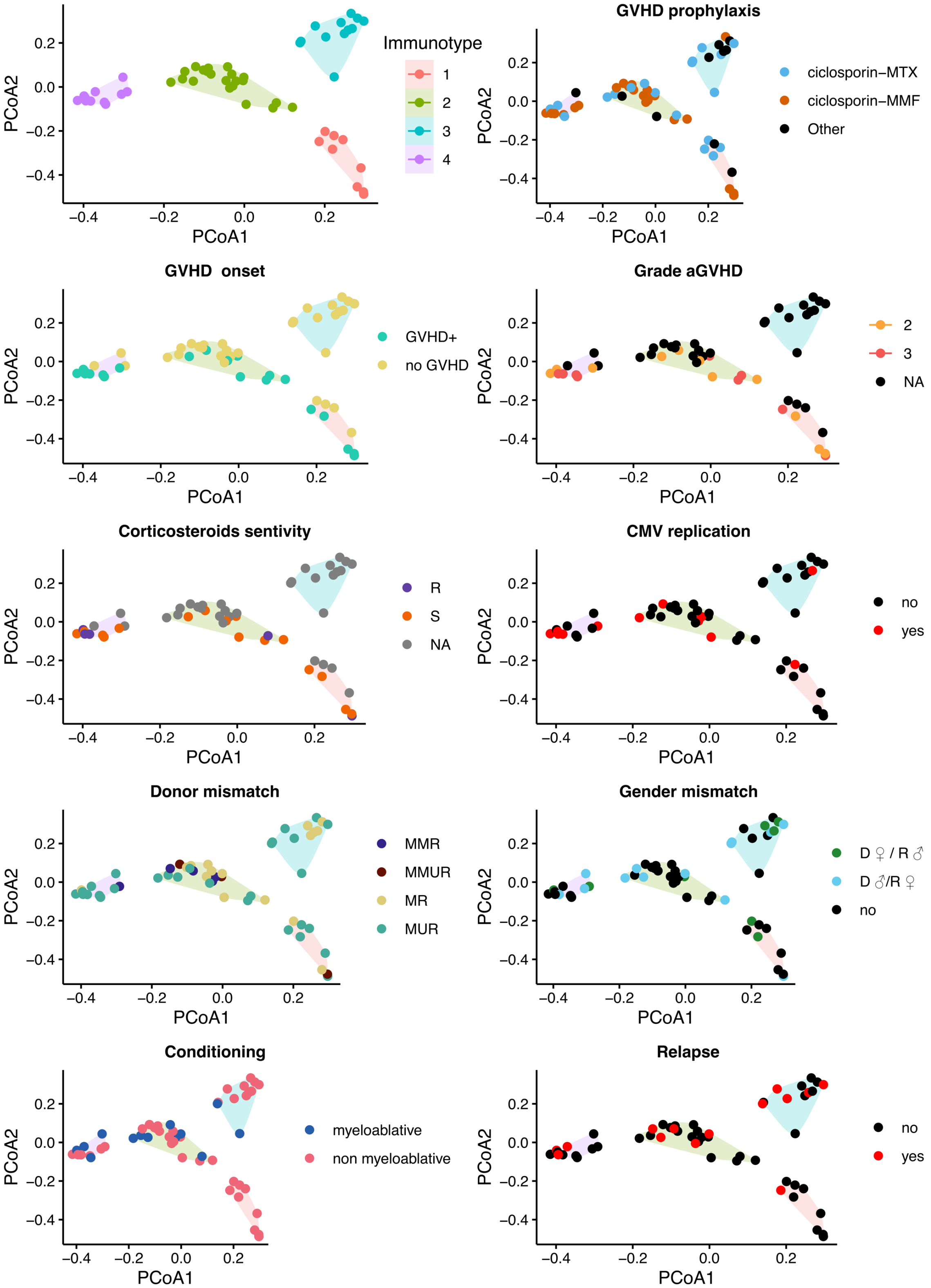


**Fig. S3: Immunotype and clinical variables:**

Clinical parameters and immunotypes.

**FIgure S4**

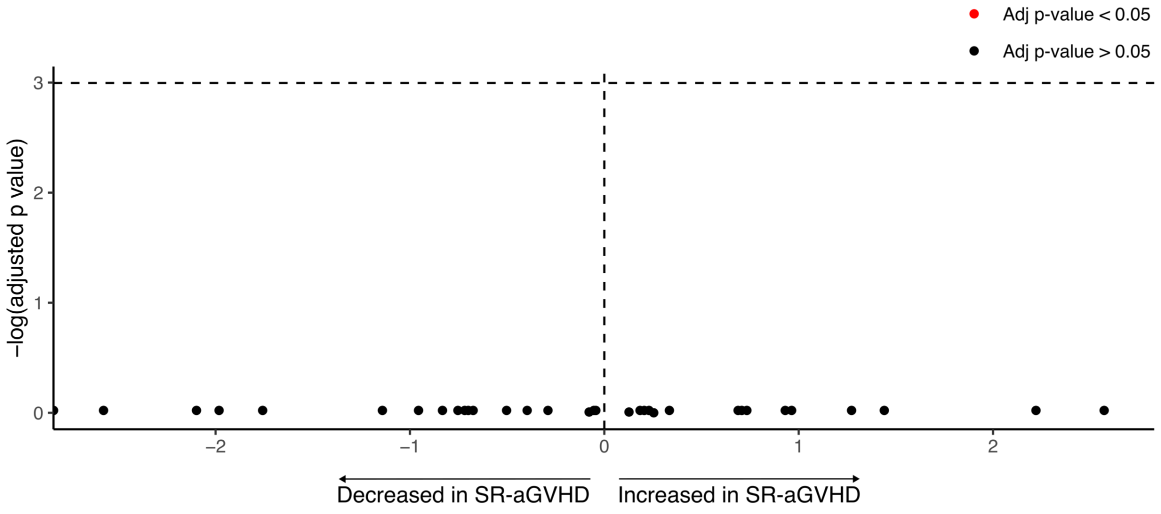


**Fig. S4: Immune cell abundance in SS- and SR-GVHD patients:**

Volcano plot exhibiting immune cells abundance in SR-GVHD and SS-GVHD condition. Y axis reflects que -log 10(adjusted p-value) from differential expression output, using non-parametric Wilcoxon rank sum test. X axis reflects the log(ratio) of median abundance in SR-GVHD condition, over SS-GVHD condition. Abbreviation: GVHD: graft versus host disease. HSCT: allogeneic hematopoietic stem cell transplantation. SS-GVHD: Steroid Sensitive acute graft versus host disease. SR-GVHD: Steroid resistant acute graft versus host disease.

**Figure S5**

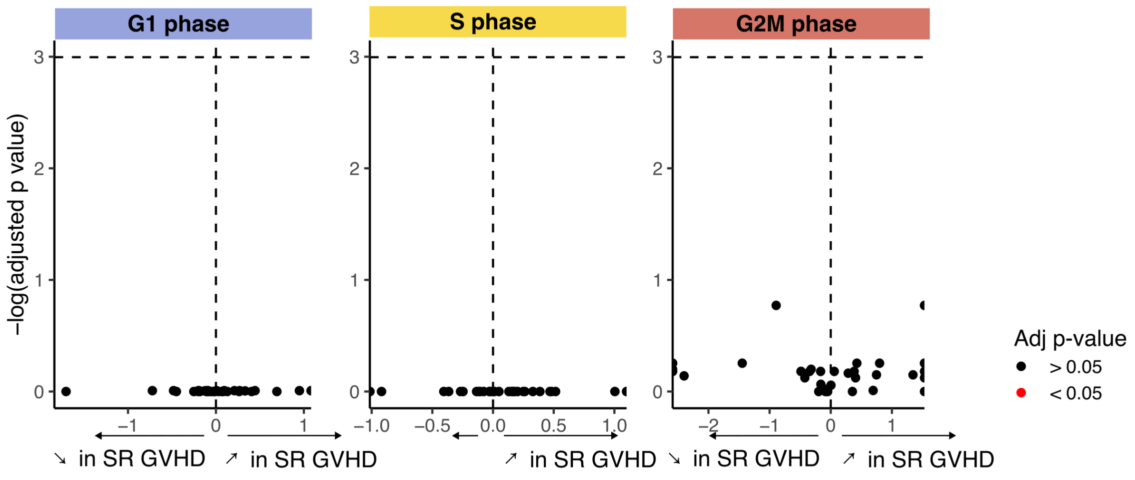


**Fig. S5: Cell cycle in SS- and SR-GVHD patients and enrichment pathways in aGVHD:** Volcano plots exhibiting cell cycle analysis after cell cycle phases scoring. X axis represents log ratio of median number of cells in each phase in SR-GVHD patients over SS-GVHD patients. Y axis represents log adjusted p values. Immune population significantly increased or decreased are marked in red. Mann-Whitney-Wilcoxon tests were performed for statistical analysis, and p values were adjusted using Benjamini and Hochberg method.

#### Figure S6

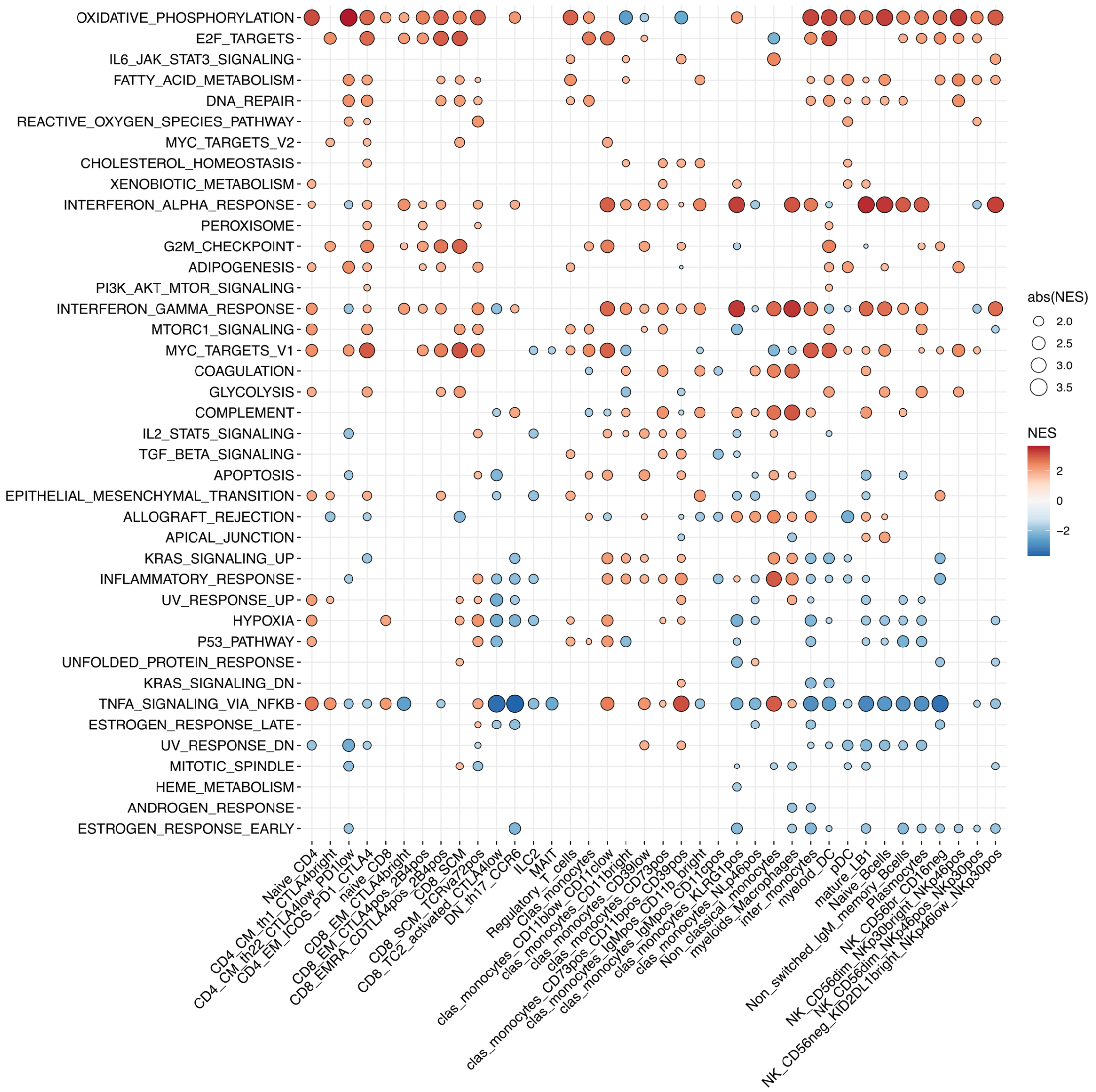


**Fig. S6: Pathway enrichments in aGVHD patients:**

Pathways enrichment analysis comparing aGVHD and no GVHD, using gene set enrichment method and "hallmarks" gene reference data set. Only significant values with adjusted p-value < 0.05 are shown. NES: normalized enrichment score. SS-GVHD: Steroid Sensitive acute graft versus host disease. SR-GVHD: Steroid resistant acute graft versus host disease.

#### Figure S7

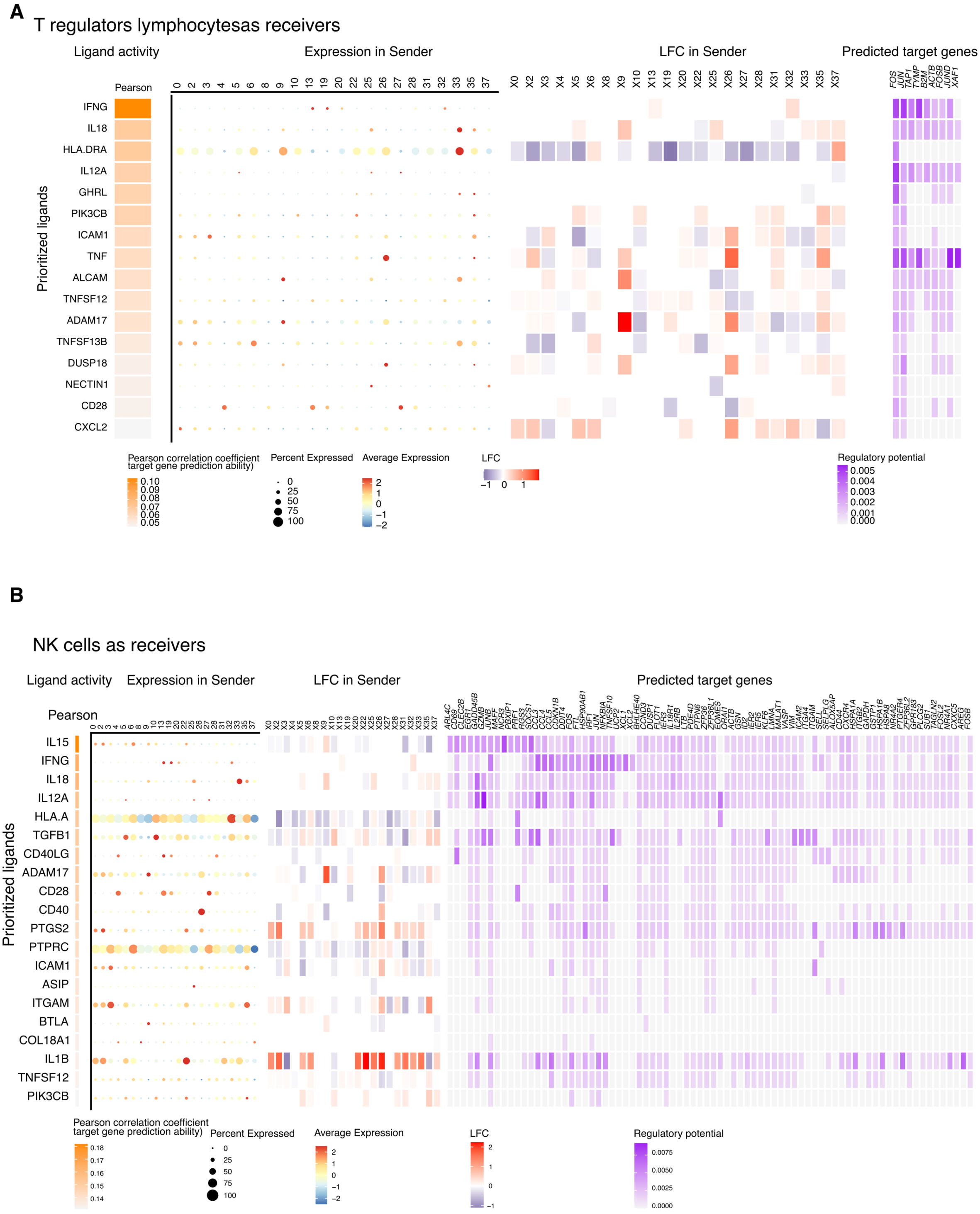


**Fig. S7: Interactions combined plots in regulatory T cells and NK cells:**

**A:** Combined plots of cell-cell interactions analysis, with T regulators lymphocytes as receiver.

**B:** Combined plots of cell-cell interactions analysis, with NK cells as receiver.

#### Figure S8

CD4$^+$ T cells as receiver

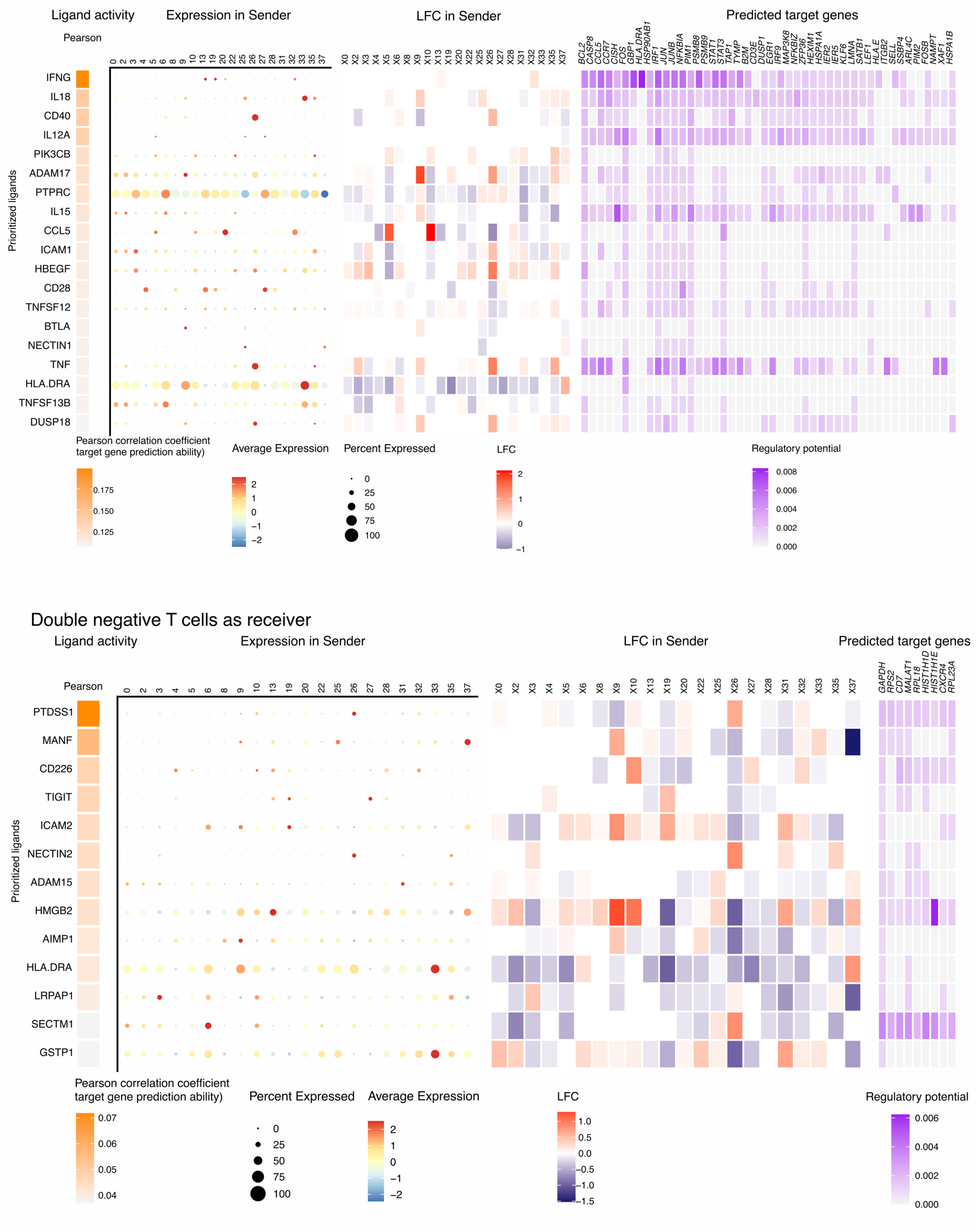


**Fig. S8: Interactions combined plots in CD4$^+$ T cells and double negative T cells**

**A:** Combined plots of cell-cell interactions analysis, with CD4$^+$ T lymphocytes as receiver.

**B:** Combined plots of cell-cell interactions analysis, with double negative T cells as receiver.

**Figure S9**

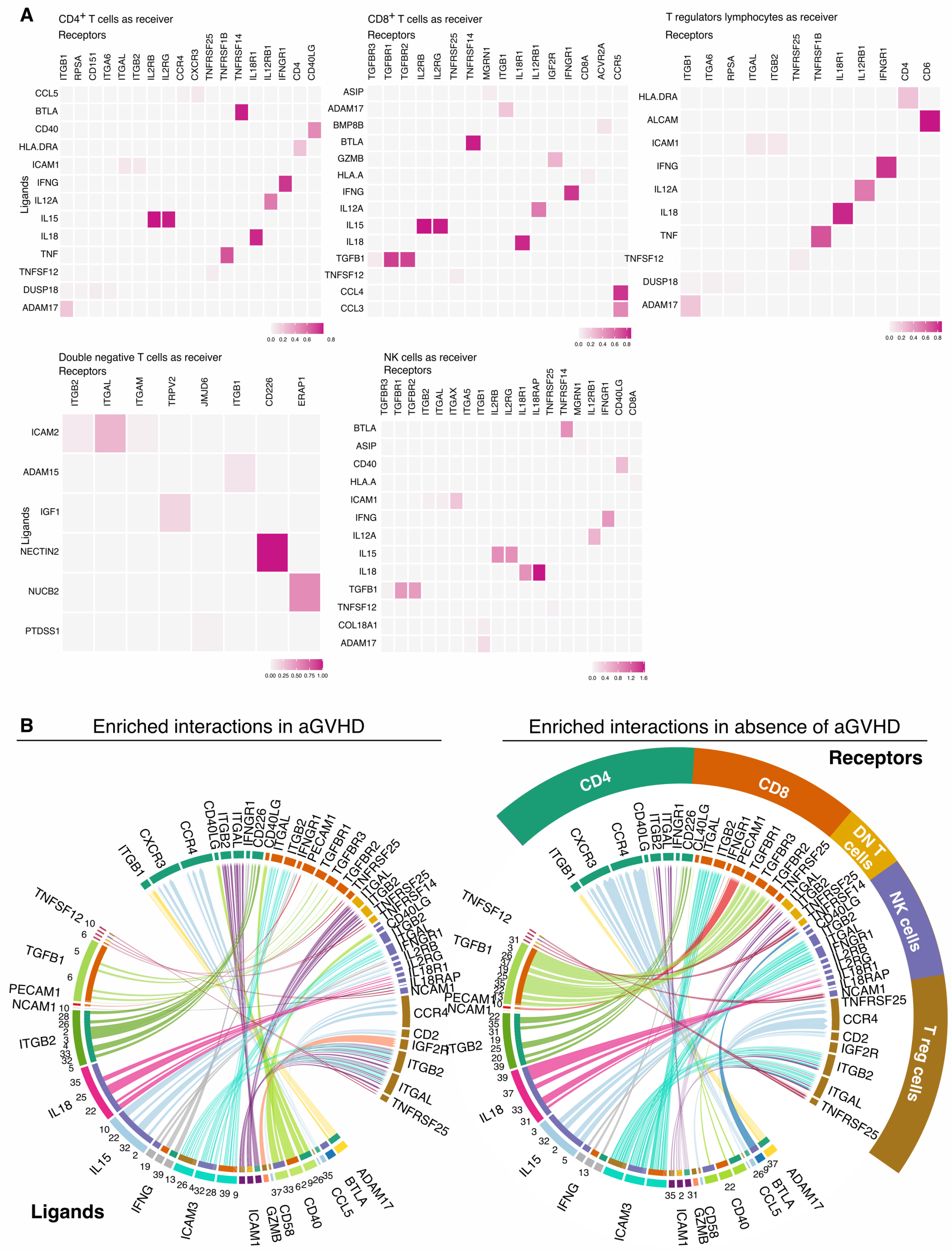


**Fig. S9: Bona fide interactions in all receivers:**

**A:** Known interactions (*bona fide*) with prioritized ligands of each receiver populations.

**B:** Increased and decreased interactions in patients with aGVHD, according to *bona fide* interactions.

#### Figure S10

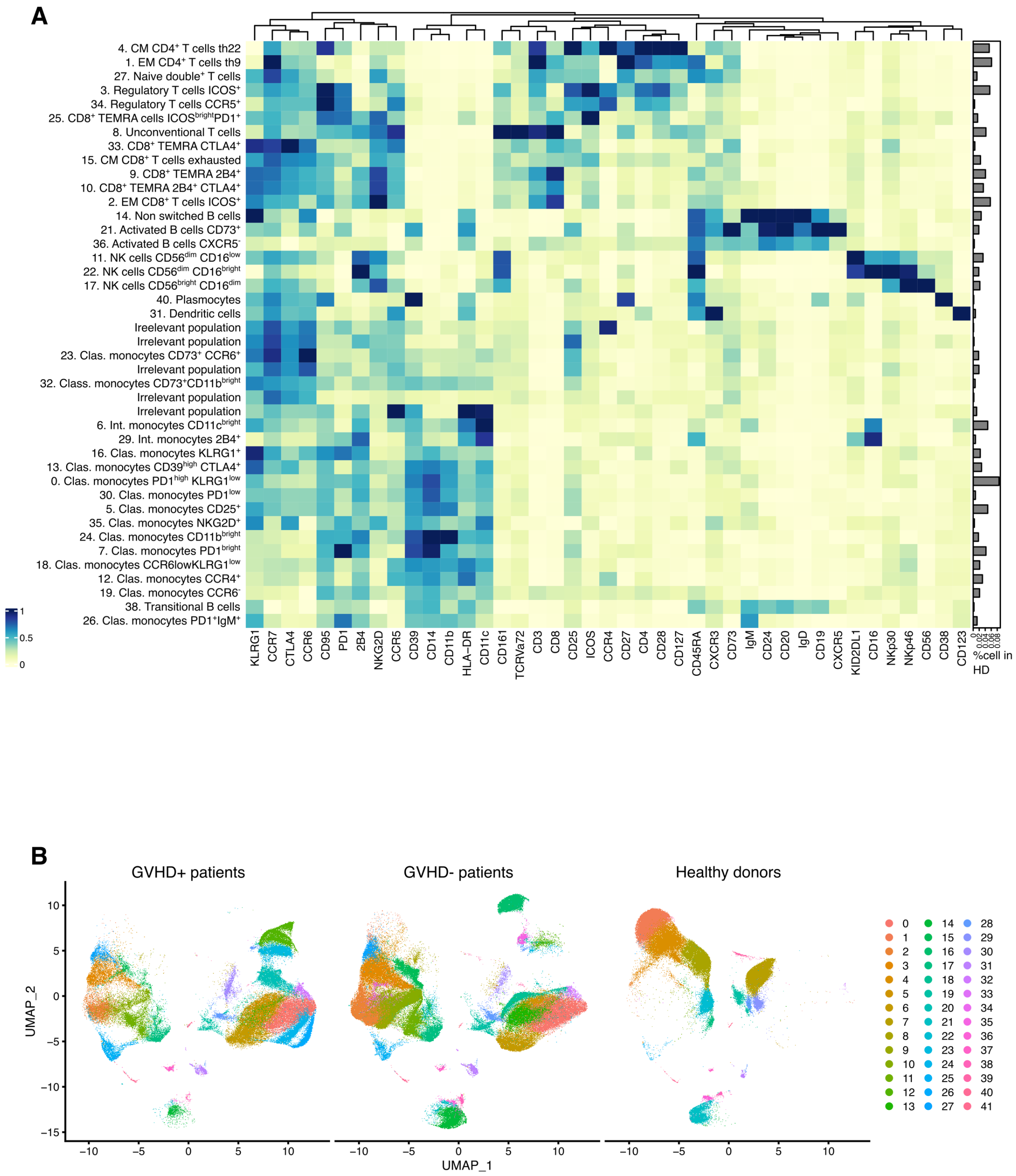


**Fig. S10: Clustering based on RNA expression**

**A:** Clusters based on mRNA expression. Immune cells were identified using protein surface expression after clustering. **B:** Immune cells on UMAP in GVHD patients, no GVHD patients and healthy donors.

**Figure S11**

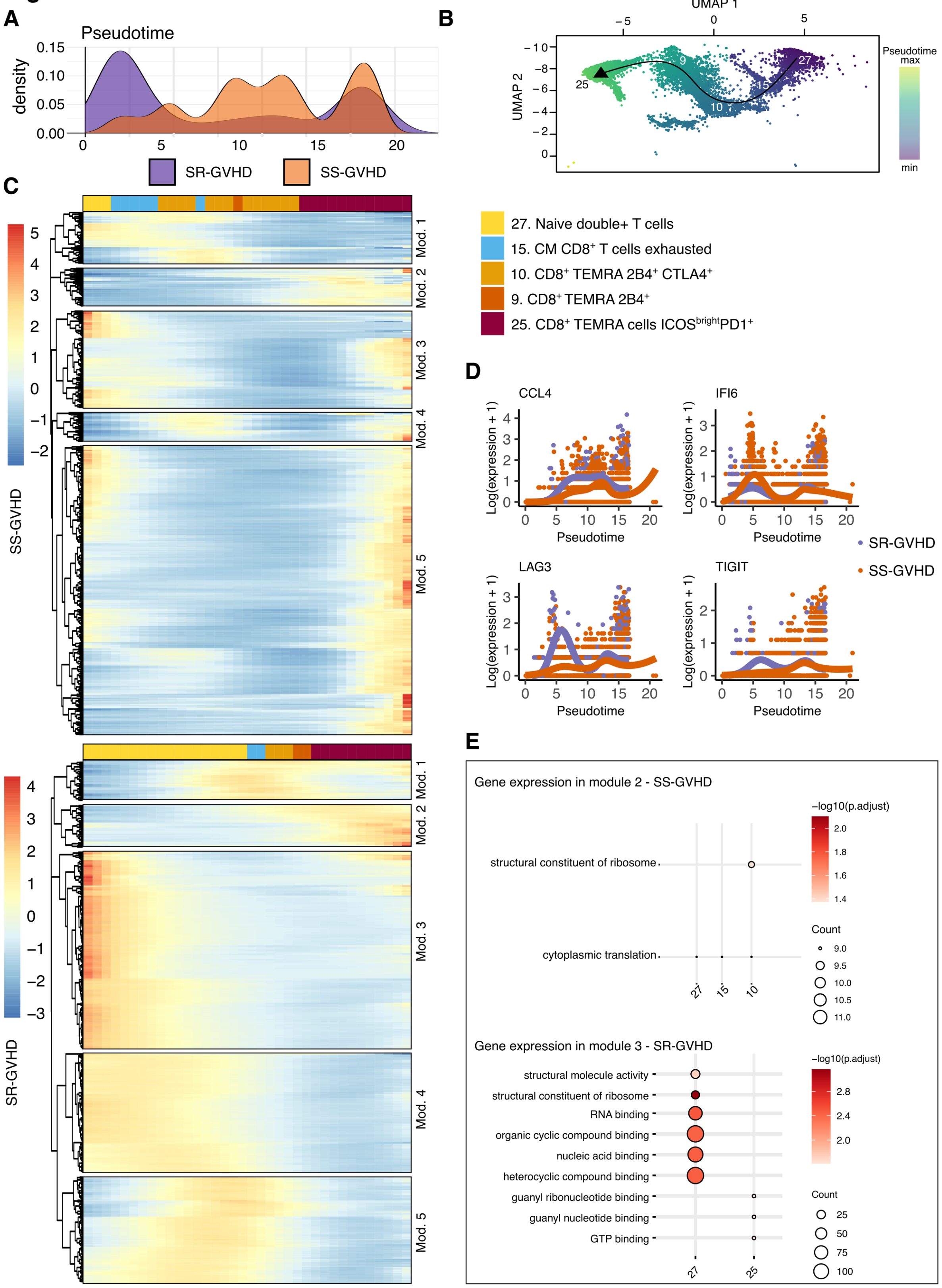


**Fig. S11: CD8$^{+}$ T cells lineage 2 trajectory analysis:**

**A:** Density plot for number of cells according to lineage 2 pseudotime visualization. SR-GVHD cell abundance was marked in low and high pseudotimes, while SS-GVHD was associated with a continuous increase in cell abundance along differentiation. **B:** Lineage 2 trajectory on RNA clustering. **C:** Heatmap of RNA expression of the 500 most expressed gene in SS-GVHD patient (top panel) and in SR-GVHD patients (low panel), along pseudotime. The color at the top of each heatmap represents the immune population in which the gene is expressed. **D:** Expression along pseudotime of four of the top 100 genes, whose expression varied the most between the start and end points of a lineage. **E:** Over-representation analysis to identify pathways associated with genes expression from each module of the heatmaps. Only pathways associated with adjusted pval < 0.05 from “molecular functions” and “biological process” of the Gene Ontology knowledgbase(*57*, *58*) were selected for dot plots visualization. SR–GVHD: Steroid resistant graft versus host disease. SS–GVHD: Steroid sensitive graft versus host disease.

### Figure S12

Lineage 1 and SR-GVHD patients

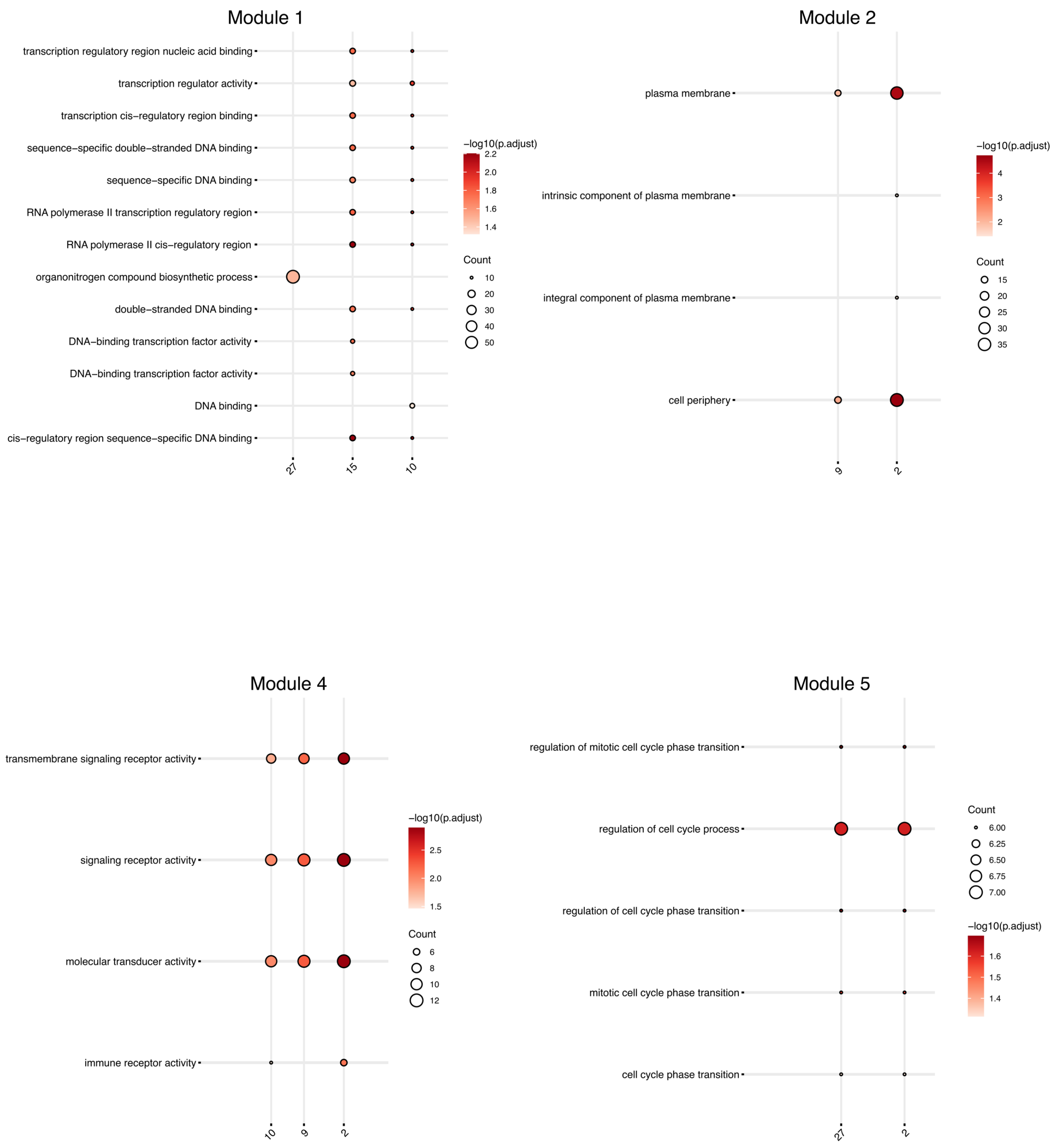


**Fig. S12: Over representation analysis in lineage 1 SR-GVHD patients:**

Over-representation analysis in lineage 1 and SR-GVHD patients to identify pathways associated with genes expression from each module of the heatmaps. Only pathways associated with adjusted pval < 0.05 from “molecular functions” and “biological process” of the Gene Ontology knowledgbase(*57*, *58*) were selected for dot plots visualization. SR–GVHD: Steroid resistant graft versus host disease. SS–GVHD: Steroid sensitive graft versus host disease.

### Figure S13

Lineage 1 and SS-GVHD patients

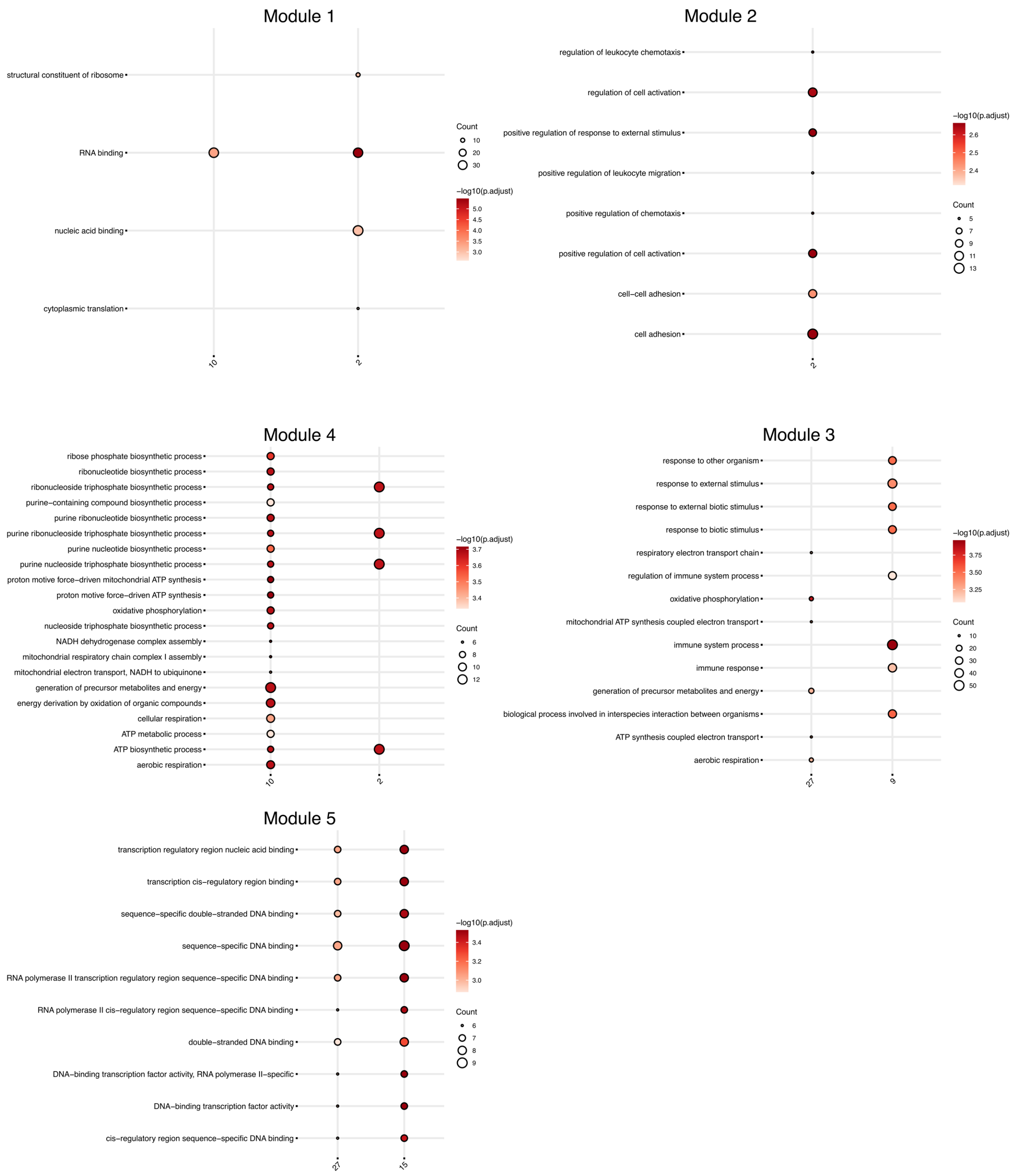


**Fig. S13: Over representation analysis in lineage 1 SS-GVHD patients:**

Over-representation analysis in lineage 1 and SS-GVHD patients to identify pathways associated with genes expression from each module of the heatmaps. Only pathways associated with adjusted pval < 0.05 from “molecular functions” and “biological process” of the Gene Ontology knowledgbase(*57*, *58*) were selected for dot plots visualization. SR–GVHD: Steroid resistant graft versus host disease. SS–GVHD: Steroid sensitive graft versus host disease.

#### Figure S14

Lineage 2 and SR-GVHD patients

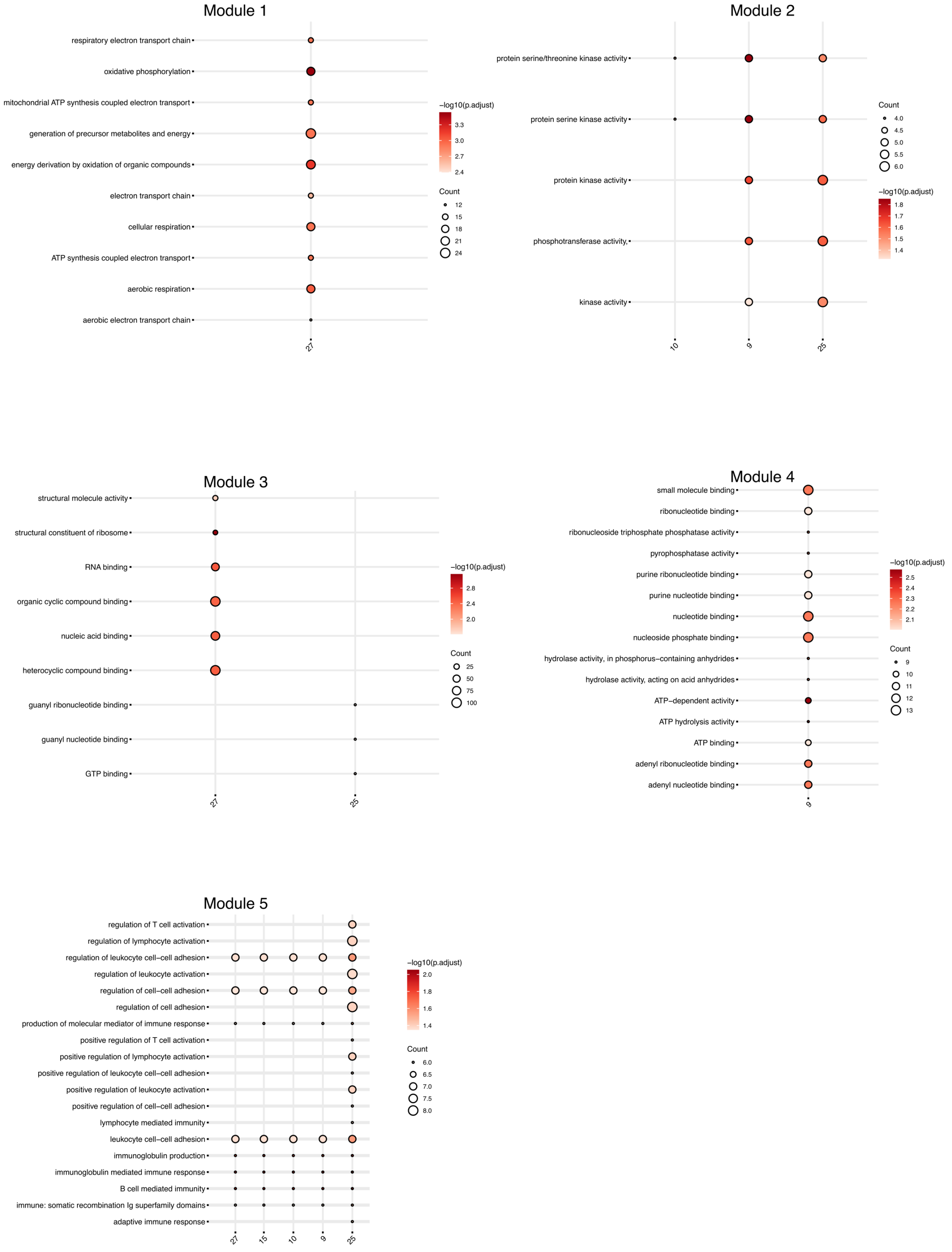


**Fig. S14: Over representation analysis in lineage 2 SR-GVHD patients:**

Over-representation analysis in lineage 2 and SR-GVHD patients to identify pathways associated with genes expression from each module of the heatmaps. Only pathways associated with adjusted pval < 0.05 from “molecular functions” and “biological process” of the Gene Ontology knowledgbase(*57*, *58*) were selected for dot plots visualization. SR–GVHD: Steroid resistant graft versus host disease. SS–GVHD: Steroid sensitive graft versus host disease.

**Figure S15**

Lineage 2 and SS GVHD patients

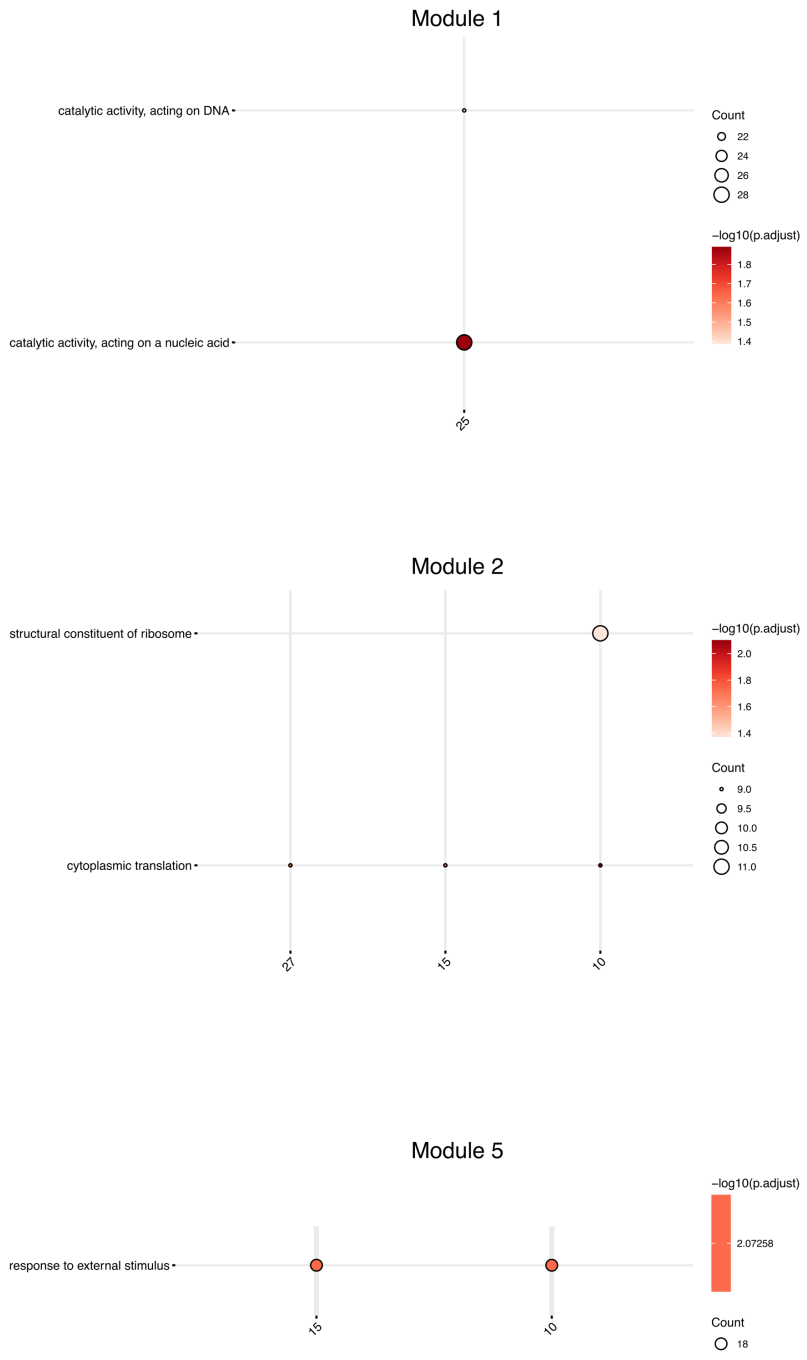


**Fig. S15: Over representation analysis in lineage 2 SS-GVHD patients:**

Over-representation analysis in lineage 2 and SS-GVHD patients to identify pathways associated with genes expression from each module of the heatmaps. Only pathways associated with adjusted pval < 0.05 from “molecular functions” and “biological process” of the Gene Ontology knowledgbase(*57*, *58*) were selected for dot plots visualization. SR–GVHD: Steroid resistant graft versus host disease. SS–GVHD: Steroid sensitive graft versus host disease.

**Supplementary tables**

**Table S1: Patients' characteristics**

| **Characteristics** | **N** | **Overall**, N = 53 | aGVHD, N = 21 | No GVHD, N = 32 | **p-value**[1] |
|---|---|---|---|---|---|
| Age, Median (years, range) | 53 | 55 (20, 69) | 57 (28, 69) | 51 (20, 69) | 0.5 |
| Sex ratio (♀/♂) | 53 | 0,47 | 0,5 | 0,45 | 0.9 |
| Diagnosis, n (%) | 53 | | | | 0.058 |
| ALL | | 4 (7.5%) | 1 (4.8%) | 3 (9.4%) | |
| AML | | 23 (43%) | 6 (29%) | 17 (53%) | |
| Biphenotypic leukemia | | 1 (1.9%) | 0 (0%) | 1 (3.1%) | |
| CML | | 1 (1.9%) | 1 (4.8%) | 0 (0%) | |
| Hodgkin lymphoma | | 2 (3.8%) | 0 (0%) | 2 (6.3%) | |
| MDS | | 13 (25%) | 8 (38%) | 5 (16%) | |
| Myelofibrosis | | 2 (3.8%) | 0 (0%) | 2 (6.3%) | |
| NHL | | 7 (13%) | 5 (24%) | 2 (6.3%) | |
| Disease status at transplantation, n (%) | 53 | | | | 0.061 |
| >CR1 | | 11 (21%) | 3 (14%) | 8 (25%) | |
| CR1 | | 20 (38%) | 5 (24%) | 15 (47%) | |
| Other | | 22 (42%) | 13 (62%) | 9 (28%) | |
| Type, n (%) | 53 | | | | 0.030 |
| bone marrow | | 6 (11%) | 0 (0%) | 6 (19%) | |
| cord blood | | 1 (1.9%) | 1 (4.8%) | 0 (0%) | |
| Peripheral blood stem cells | | 46 (87%) | 20 (95%) | 26 (81%) | |
| Conditioning, n (%) | 53 | | | | 0.7 |
| myeloablative | | 14 (26%) | 5 (24%) | 9 (28%) | |
| non myeloablative | | 39 (74%) | 16 (76%) | 23 (72%) | |
| Donor, n (%) | 53 | | | | 0.4 |
| Matched related | | 16 (30%) | 4 (19%) | 12 (38%) | |

| **Characteristics** | **N** | **Overall**, N = 53 | aGVHD, N = 21 | No GVHD, N = 32 | **p-value**[1] |
|---|---|---|---|---|---|
| Matched unrelated | | 29 (55%) | 13 (62%) | 16 (50%) | |
| Haploidentical | | 5 (9.4%) | 2 (9.5%) | 3 (9.4%) | |
| Mismatched unrelated | | 3 (5.7%) | 2 (9.5%) | 1 (3.1%) | |
| GVHD prophylaxis, n (%) | 53 | | | | 0.093 |
| cyclosporin-MTX | | 20 (38%) | 6 (29%) | 14 (44%) | |
| cyclosporin-MMF | | 23 (43%) | 13 (62%) | 10 (31%) | |
| Other | | 10 (19%) | 2 (9.5%) | 8 (25%) | |
| Grade GVHD, n (%) | 53 | | | | |
| 2 | | 12 (23%) | 12 (57%) | 0 (0%) | |
| 3 | | 9 (17%) | 9 (43%) | 0 (0%) | |
| 0 | | 32 (60%) | 0 (0%) | 32 (100%) | |
| GVHD organ involvement | | | | | |
| Skin | | 19 (36%) | 19 (90%) | NA | |
| Gut | | 7 (13%) | 7 (33%) | NA | |
| Time between transplantation and sample (days), Median (Range) | 53 | 84 (11-106) | 21 (11, 96) | 89 (75,106) | <0.001 |
| Steroid response, n (%) | 21 | | | | |
| Resistance | | 5 (24%) | 5 (24%) | NA | |
| Sensitive | | 16 (76%) | 16 (76%) | NA | |
| Relapse, n (%) | 53 | 14 (26%) | 4 (19%) | 10 (31%) | 0.3 |
| Delay relapse from HSCT, Median (Range) | 53 | 23 (3, 37) | 24 (3, 26) | 22 (4, 37) | >0.9 |
| CMV D/R status, n (%) | 53 | | | | 0.2 |
| D-/R- | | 19 (36%) | 11 (52%) | 8 (25%) | |

| Characteristics | N | **Overall**, N = 53 | aGVHD, N = 21 | No GVHD, N = 32 | **p-value**[1] |
|---|---|---|---|---|---|
| D-/R+ | | 12 (23%) | 3 (14%) | 9 (28%) | |
| D+/R- | | 5 (9.4%) | 1 (4.8%) | 4 (13%) | |
| D+/R+ | | 17 (32%) | 6 (29%) | 11 (34%) | |
| CMV reactivation, n (%) | 53 | 12 (23%) | 6 (29%) | 6 (19%) | 0.5 |
| ABO mismatch, n (%) | 53 | | | | 0.7 |
| Major incompatibility | | 10 (19%) | 5 (24%) | 5 (16%) | |
| Minor incompatibility | | 14 (26%) | 6 (29%) | 8 (25%) | |
| No mismatch | | 29 (55%) | 10 (48%) | 19 (59%) | |
| Gender mismatch, n (%) | 53 | | | | >0.9 |
| D ♀ / R ♂ | | 8 (15%) | 3 (14%) | 5 (16%) | |
| D ♂/R ♀ | | 13 (25%) | 5 (24%) | 8 (25%) | |
| No mismatch | | 32 (60%) | 13 (62%) | 19 (59%) | |
| Follow up from HSCT, Median (years, range) | 53 | 6.83 (0.40, 7.77) | 7.02 (0.40, 7.77) | 6.78 (0.47, 7.69) | 0.3 |

[1]Wilcoxon rank sum exact test; Pearson's Chi-squared test; Fisher's exact test; Wilcoxon rank sum test

**Table S2: Characteristics according to immunotypes**

| Characteristic | N | Overall<br>N = 53[1] | Immunotype 1<br>N = 9[1] | Immunotype 2<br>N = 21[1] | Immunotype 3<br>N = 12[1] | Immunotype 4<br>N = 11[1] | p-value[2] |
|---|---|---|---|---|---|---|---|
| Group, n(%) | 53 | | | | | | 0.001 |
| aGVHD+ | | 21 (40%) | 5 (56%) | 8 (38%) | 0 (0%) | 8 (73%) | |
| No GVHD | | 32 (60%) | 4 (44%) | 13 (62%) | 12 (100%) | 3 (27%) | |
| Steroid response | 21 | | | | | | 0.8 |
| Resistance | | 5 (24%) | 1 (20%) | 1 (13%) | 0 (NA%) | 3 (38%) | |
| Sensitive | | 16 (76%) | 4 (80%) | 7 (88%) | 0 (NA%) | 5 (63%) | |
| GVHD organ involvement | | | | | | | 0.2 |
| Skin | | 19 (36%) | 5 (56%) | 6 (29%) | 0 (NA) | 8 (73%) | |
| Gut | | 7 (13%) | 0 (0%) | 5 (24%) | 0 (NA) | 2 (18%) | |
| Age (at transplantation), med(min-max) | 53 | 55(20 - 69) | 64(48 - 68) | 49(27 - 69) | 58(29 - 69) | 55(20 - 67) | 0.033 |
| Sex ratio (male/female) | 53 | 2.1 | 3.5 | 2 | 1.4 | 2.7 | 0.8 |
| Gender mismatch | 53 | | | | | | 0.5 |
| ♀/♂ | | 8 (15%) | 2 (22%) | 1 (4.8%) | 3 (25%) | 2 (18%) | |
| ♂/♀ | | 13 (25%) | 1 (11%) | 5 (24%) | 4 (33%) | 3 (27%) | |
| No mismatch | | 32 (60%) | 6 (67%) | 15 (71%) | 5 (42%) | 6 (55%) | |
| Hematological malignancy | 53 | | | | | | 0.3 |
| Acute lymphoid leukemia | | 4 (7.5%) | 0 (0%) | 1 (4.8%) | 2 (17%) | 1 (9.1%) | |
| Acute myeloid leukemia | | 23 (43%) | 4 (44%) | 10 (48%) | 7 (58%) | 2 (18%) | |
| Biphenotypic leukemia | | 1 (1.9%) | 0 (0%) | 1 (4.8%) | 0 (0%) | 0 (0%) | |

| Characteristic | N | Overall<br>N = 53[1] | Immunotype 1<br>N = 9[1] | Immunotype 2<br>N = 21[1] | Immunotype 3<br>N = 12[1] | Immunotype 4<br>N = 11[1] | p-value[2] |
|---|---|---|---|---|---|---|---|
| Chronic myeloid leukemia | | 1 (1.9%) | 0 (0%) | 0 (0%) | 0 (0%) | 1 (9.1%) | |
| Hodgkin lymphoma | | 2 (3.8%) | 0 (0%) | 0 (0%) | 1 (8.3%) | 1 (9.1%) | |
| Myelodysplastic syndrome | | 13 (25%) | 3 (33%) | 5 (24%) | 1 (8.3%) | 4 (36%) | |
| Myelofibrosis | | 2 (3.8%) | 1 (11%) | 0 (0%) | 1 (8.3%) | 0 (0%) | |
| Non Hodgkin lymphoma | | 7 (13%) | 1 (11%) | 4 (19%) | 0 (0%) | 2 (18%) | |
| Pre-HSCT disease status, n(%) | 53 | | | | | | 0.6 |
| >CR1 | | 11 (21%) | 1 (11%) | 4 (19%) | 5 (42%) | 1 (9.1%) | |
| CR1 | | 20 (38%) | 4 (44%) | 8 (38%) | 4 (33%) | 4 (36%) | |
| NA | | 22 (42%) | 4 (44%) | 9 (43%) | 3 (25%) | 6 (55%) | |
| HSCT origin of cells, n(%) | 53 | | | | | | 0.3 |
| Bone marrow | | 6 (11%) | 0 (0%) | 3 (14%) | 3 (25%) | 0 (0%) | |
| Cord blood | | 1 (1.9%) | 0 (0%) | 1 (4.8%) | 0 (0%) | 0 (0%) | |
| Peripheral blood stem cells | | 46 (87%) | 9 (100%) | 17 (81%) | 9 (75%) | 11 (100%) | |
| GVHD prophylaxis, n(%) | 53 | | | | | | 0.056 |
| Ciclosporin-MTX | | 20 (38%) | 4 (44%) | 6 (29%) | 6 (50%) | 4 (36%) | |
| Ciclosporin-MMF | | 23 (43%) | 3 (33%) | 13 (62%) | 1 (8.3%) | 6 (55%) | |
| Other | | 10 (19%) | 2 (22%) | 2 (9.5%) | 5 (42%) | 1 (9.1%) | |
| Donor type, n(%) | 53 | | | | | | 0.2 |
| Haploidentical | | 5 (9.4%) | 0 (0%) | 4 (19%) | 0 (0%) | 1 (9.1%) | |

| **Characteristic** | **N** | **Overall**<br>N = 53[1] | **Immunotype 1**<br>N = 9[1] | **Immunotype 2**<br>N = 21[1] | **Immunotype 3**<br>N = 12[1] | **Immunotype 4**<br>N = 11[1] | **p-value**[2] |
|---|---|---|---|---|---|---|---|
| MMUD | | 3 (5.7%) | 1 (11%) | 2 (9.5%) | 0 (0%) | 0 (0%) | |
| MRD | | 16 (30%) | 2 (22%) | 8 (38%) | 5 (42%) | 1 (9.1%) | |
| MUD | | 29 (55%) | 6 (67%) | 7 (33%) | 7 (58%) | 9 (82%) | |
| Conditioning | 53 | | | | | | 0.090 |
| Myeloablative | | 14 (26%) | 0 (0%) | 7 (33%) | 2 (17%) | 5 (45%) | |
| Non myeloablative | | 39 (74%) | 9 (100%) | 14 (67%) | 10 (83%) | 6 (55%) | |
| ABO incompatibility, n(%) | 53 | | | | | | 0.7 |
| Major | | 10 (19%) | 3 (33%) | 4 (19%) | 2 (17%) | 1 (9.1%) | |
| Minor | | 14 (26%) | 3 (33%) | 4 (19%) | 3 (25%) | 4 (36%) | |
| No incompatibility | | 29 (55%) | 3 (33%) | 13 (62%) | 7 (58%) | 6 (55%) | |
| Donor/recipient CMV status, n(%) | 53 | | | | | | >0.9 |
| D-/R- | | 19 (36%) | 3 (33%) | 6 (29%) | 4 (33%) | 6 (55%) | |
| D-/R+ | | 12 (23%) | 2 (22%) | 4 (19%) | 3 (25%) | 3 (27%) | |
| D+/R- | | 5 (9.4%) | 1 (11%) | 3 (14%) | 1 (8.3%) | 0 (0%) | |
| D+/R+ | | 17 (32%) | 3 (33%) | 8 (38%) | 4 (33%) | 2 (18%) | |
| CMV replication, n(%) | 53 | 12 (23%) | 1 (11%) | 5 (24%) | 1 (8.3%) | 5 (45%) | 0.2 |
| Relapse, n(%) | 53 | 14 (26%) | 1 (11%) | 5 (24%) | 5 (42%) | 3 (27%) | 0.5 |

[1]n (%); Median(Range)

[2]Fisher's exact test; Kruskal-Wallis rank sum test

114

**Supplementary data files**

Data file S3: Top 500 genes expression matrix in lineage 1 SR-GVHD patients

Data file S4: Top 500 genes expression matrix in lineage 1 SS-GVHD patients

Data file S5: Top 500 genes expression matrix in lineage 2 SR-GVHD patients

Data file S6: Top 500 genes expression matrix in lineage 2 SS-GVHD patients

Data file S7: Immune population mostly represented along pseudotimes in lineage 1 SR-GVHD patients

Data file S8: Immune population mostly represented along pseudotimes in lineage 1 SS-GVHD patients

Data file S9: Immune population mostly represented along pseudotimes in lineage 1 2 SR-GVHD patients

Data file S10: Immune population mostly represented along pseudotimes in lineage 2 SS-GVHD patients

Data file S11: StartvsEnd output in lineage 1

Data file S12: StartvsEnd output in lineage 2

Data file S13: Total-Seq-B antibody panel
